# Characterization of a *RAD51C*-Silenced High Grade Serous Ovarian Cancer Model During PARP Inhibitor Resistance Development

**DOI:** 10.1101/2020.12.25.419978

**Authors:** Rachel M. Hurley, Cordelia D. McGehee, Ksenija Nesic, Cristina Correia, Taylor M. Weiskittel, Rebecca L. Kelly, Annapoorna Venkatachalam, Xiaonan Hou, Nicholas M. Pathoulas, X. Wei Meng, Olga Kondrashova, Marc R. Radke, Paula A. Schneider, Karen S. Flatten, Kevin L. Peterson, Alexander Dobrovic, Kevin K. Lin, Thomas C Harding, Iain A. McNeish, Christian A. Ross, Jill M. Wagner, Matthew J. Wakefield, Clare L. Scott, Paul Haluska, Andrea E. Wahner Hendrickson, Larry M. Karnitz, Elizabeth M. Swisher, Hu Li, S. John Weroha, Scott H. Kaufmann

## Abstract

Acquired PARP inhibitor (PARPi) resistance in *BRCA1*- or *BRCA2*-mutant ovarian cancer often results from secondary mutations that restore expression of functional protein. *RAD51C* is a less commonly studied ovarian cancer susceptibility gene whose promoter is sometimes methylated in the tumor, leading to homologous recombination deficiency and PARPi sensitivity. For this study, the PARPi-sensitive patient-derived xenograft PH039, which lacks demonstrable repair gene mutations but harbors *RAD51C* promoter methylation, was selected for PARPi resistance by repeated 21-day niraparib treatments *in vivo*. PH039 acquired PARPi resistance by the third cycle of treatment and demonstrated unimpeded growth during subsequent exposure to either niraparib or rucaparib. Transcriptional profiling throughout the time course of resistance development showed widespread pathway level changes along with a marked increase in *RAD51C* mRNA, which reflected loss of *RAD51C* promoter methylation. Analysis of *RAD51C* methylation in patient tumor samples from the ARIEL2 Part 1 clinical trial of rucaparib monotherapy likewise indicated that loss of *RAD51C* methylation prior to on-study biopsy was associated with limited response. Interestingly, the PARPi resistant PH039 model remained platinum sensitive. Collectively, these results not only indicate that PARPi treatment pressure can reverse *RAD51C* methylation and restore RAD51C expression, but also provide an important model for studying the clinical observation that PARPi and platinum sensitivity are sometimes dissociated.

## INTRODUCTION

Up to 50% of high-grade serous ovarian carcinomas (HGSOC) display homologous recombination (HR) deficiency (1,2). The lesions most commonly implicated are mutations in *BRCA1* and *BRCA2*, which are present in 17-25% of HGSOCs (1–6). An additional 10-15% of HGSOC cases have lost BRCA1 expression as a consequence of CDK12 inactivation (1,7,8) or *BRCA1* promoter methylation (1,2,9–11), which often reflects loss of the zinc finger protein ZC3H18 (12). Whatever the cause, the resulting loss of BRCA1 or BRCA2 is associated with increased sensitivity to PARP inhibitors (PARPis) (13–15), which are approved for ovarian cancer both as maintenance therapy and as monotherapy for relapsed, platinum-sensitive disease (16–18).

The lesions contributing to BRCA1 or BRCA2 loss are often reversed when HGSOCs lacking these proteins become therapy resistant (19). For example, secondary mutations that restore the *BRCA1* or *BRCA2* open reading frame have been observed in DNA from platinum- (20) and PARPi-resistant (15,21,22) HGSOC. Additionally, loss of promoter methylation in a single copy of the *BRCA1* gene was shown to be sufficient to restore HR and cause PARPi resistance (23).

Another 6-10% of HGSOC cases harbor mutations in other genes that contribute to HR (1,3–6,24–26). These inactivated genes include *RAD51C* and *RAD51D*, which encode RAD51 paralogs that participate in HR repair by modulating the action of the RAD51 recombinase (27). Previous studies have established that HGSOCs with *RAD51C* and *RAD51D* mutations are sensitive to the PARPi rucaparib (25). Conversely, secondary mutations that restore the *RAD51C* or *RAD51D* open reading frame and reestablish HR are observed when *RAD51C*- or *RAD51D*-mutant HGSOCs become rucaparib resistant (15,25). Promoter hypermethylation resulting in silencing of *RAD51C* has also been reported in 1-2% of ovarian cancers (1,2,9–11). Less is currently known about the changes that lead to PARPi resistance in *RAD51C* methylated tumors.

HR alterations that modulate PARPi sensitivity generally affect sensitivity to platinating agents as well. In preclinical models, loss of HR components sensitizes to both drug classes and restoration of HR produces resistance to both (23,25,28,29). Moreover, the degree of ovarian cancer shrinkage with the PARPi olaparib in the clinical setting has been correlated with platinum-free interval (an index of platinum sensitivity) (30). Despite this general correlation, olaparib responses at any particular platinum-free interval are quite variable (30), raising the possibility that factors other than HR deficiency might also play important roles in platinum sensitivity. Consistent with this notion, defects in nucleotide excision repair have been reported to sensitize HGSOCs to cisplatin without affecting PARPi sensitivity (31). It has also been observed that 40% of ovarian cancers progressing on PARPi treatment will respond to subsequent platinum-based therapy (32), although the basis for these divergent responses remains unclear.

Patient-derived xenograft (PDX) models can provide a unique view into acquired resistance. In particular, intraperitoneal HGSOC PDX models retain critical features, including the pathogenic mutations, copy number alterations, gene expression profiles, degree of stromal infiltration, propensity to metastasize to bowel or induce ascites, sensitivity or resistance to drug treatment, and tumor heterogeneity, of the original cancers (33–35), providing an opportunity for studying drug resistance under conditions somewhat similar to the original cancers. While acquired drug resistance studies in solid tumor PDX models typically rely on endpoint analyses, losing much of the time-course resolution of changes, serial passaging of PDX models can also allow for time resolution of signaling changes while simultaneously controlling for tumor evolution in a matched untreated control.

At the present time little is known about the stability of *RAD51C* methylation under PARPi treatment pressure. In the clinical setting, it is also difficult to determine whether alterations that affect PARPi sensitivity concomitantly affect platinum sensitivity. Because sister tumors can be simultaneously assayed for response to multiple agents, PDXs are a potentially important tool for addressing these questions. Here we report the loss of *RAD51C* methylation over a time course of acquired PARPi resistance in a PDX model and its differential impact on PARPi versus platinum sensitivity.

## METHODS

### Reagents

Rucaparib was provided by Clovis Oncology. Antibodies for immunoblotting were purchased as follows: Murine anti-RAD51C (NB100-177) from Novus (Centennial, CO), murine anti-RAD51 (MS-988-P0) from Thermo (Fremont, CA) and murine anti-LMNB (sc-377000) from Santa Cruz Biotechnology. Chicken polyclonal anti-B23/nucleophosmin was raised as previously described (36).

### Resistance Development in Vivo

PDX PH039 was established intraperitoneally utilizing previously described methods (33). Briefly, tissue was transplanted into the peritoneal cavities of five female SCID Beige mice (C.B-17/IcrHsd-*Prkdc^scid^Lyst^bg-^*J; Envigo, Indianapolis, IN) and tumor size (largest cross-sectional area) was monitored by transabdominal ultrasound (SonoSite S-Series Ultrasound, Fujifilm SonoSite, Bothell, WA). At a tumor area of 0.5-0.8 cm^2^, mice were randomized to the niraparib arm (n=3) or source tumor arm (n=2) as indicated in **Fig. 1A**. On the source tumor arm, tumors were monitored for growth, harvested, and re-established in two mice. Tumor was passaged five times to control for any changes unrelated to niraparib treatment. On the niraparib arm, mice treated with 100 mg/kg niraparib (dissolved in 0.5% methylcellulose) daily for 21 days via oral gavage were monitored for regrowth of tumors, which were harvested and re-established in three mice. Once established, the tumor was again treated with niraparib for 21 days. This continued until the tumor grew through treatment in two sequential passages. Tumor size was monitored twice weekly, and mice were euthanized when moribund or followed for time to tumor regrowth, in accordance with Mayo Clinic Institutional Animal Care and Use Committee protocols. Additional details of the animal studies are described in the Supplemental Methods.

**Figure 1.**
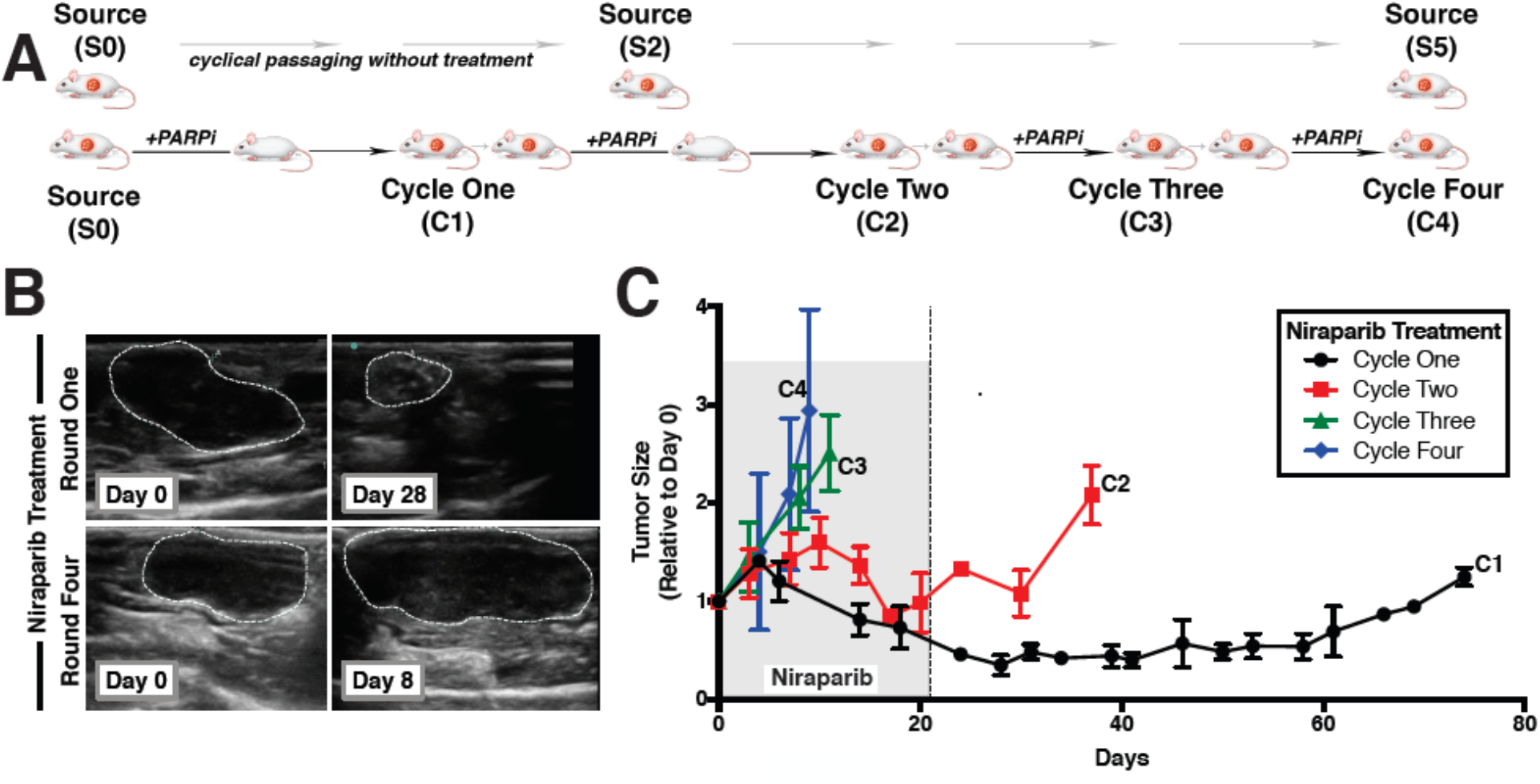
Development of PARPi resistance in PH039. **A,** to develop PARP inhibitor resistance, PH039, an intraperitoneal ovarian cancer PDX with no known mutations in HR, was treated with niraparib (100 mg/kg) daily via oral gavage for 21 days. Tumors were subsequently allowed to regrow and were then re-established in 3 new mice. This process was repeated for a total of four treatment cycles of niraparib. Shaded area, period of niraparib treatment. Tumor was also passaged five times without treatment to account for any genetic drift with passaging alone. **B,** representative ultrasounds showing shrinkage of PH039 with first treatment cycle and lack of shrinkage with fourth treatment cycle. **C,** tumor growth throughout the study was assessed by ultrasound.

### RNA sequencing

RNA was harvested from snap-frozen xenograft tissue using RNeasy Plus Minikit (Qiagen, Germany) or a Direct-zol RNA MiniPrep (Zymo Research, Cat# R2050). RNA sample quality was assessed by RNA integration number (RIN) on the RNA ScreenTape System (Agilent) according to manufacturer’s protocol. cDNA was generated with the Illumina TruSeq mRNA kit. The cDNAs were then denatured and polymerase chain reaction (PCR) enriched. The resulting genomic library was analyzed on an Illumina HiSeq 4000.

### Bioinformatic Analysis of RNAseq Data

The bioinformatic pipeline is outlined in **Figure S1**. Tophat (37,38) was used for sequence alignment to the UCSC HG38 (human) or UCSC MM10 (mouse) in conjunction with Picard (http://broadinstitute.github.io/picard). (See Supplementary Material for description and validation of method for addressing multi-mapping of sequences.) Aligned reads were quantified using Subread v1.4.6 (39,40). Differential expression analysis was performed with EdgeR (41) and Bioconductor v3.4 (42) in R (43). For pairwise comparisons, gene filtering was set to include any gene with a total of more than two or three counts per million (two if two was the lowest number of samples for a given condition in the comparison and three if three was the lowest number of samples for a given condition.). IPA pathway analysis (44) (QIAGEN Inc., https://www.qiagenbioinformatics.com/products/ingenuitypathway-analysis) was performed for genes with |log2 FC | > 1 and p-value< 0.01 unless otherwise noted. Figures were generated in R using ComplexHeatmap (45), pvclust (46), dendextend (47), MASS (48), circlize (49), colorspace (50), GetoptLong (51), and ggplot2 (52). Heatmap colors based on www.ColorBrewer.org by Cynthia A. Brewer, Penn State. Genes corresponding to the ontology tag DNA Repair (GO:0006281) in Homo Sapiens were downloaded from AmiGO 2 (53,54) and intersected with differentially expressed genes.

### SNV Calling

SAMtools (v1.3.1) (55) was applied to BAM aligned and mouse depleted files to generate mpileups against GRCh38.86 human reference genome. VarScan2 (v2.4.1) (56) was used for baseline calling (S0 samples #1 and #2 against the human genome). Somatic calling using S0 as reference was performed for two samples passaged without selection (S2 #1, S2 #2) and two niraparib-resistant samples (C3 #1 and C3 #2). Reads were filtered for min-coverage of 10 and min-base-qual of 20. This allowed us to identify common variants at each time point that may contribute to the observed cisplatin sensitive phenotype. Variants were annotated with SNPEFFECT (57) and dbSNP15 was used for SNP annotations. For SNV selection, we filtered high confidence missense somatic variants and indels (p-value somatic <0.05). We inspected variants for 281 DNA damage repair genes (58), and all ATP transporters (*ATP) and solute carriers (*SLC), including all (SLC31 family of copper transporters (CTR1, CTR2) and ATOX1 and CCS) previously described to impact cisplatin sensitivity (59).

### Transcription Factor Analysis

Transcription factor analysis was performed focusing on transcription factors and targets that are expressed in the PDX samples. Log2 average fold change across all samples was compared to S0, and C3 was compared to S2 for each gene. Transcription factors, targets, and their mode of regulation (“Activation”, “Repression”, “Unknown”) were extracted from the TRRUST v2 database (60). Entries with “Unknown” modes of regulation and transcription factors with less than five annotation entries were excluded from the analysis. A transcription factor activity score was calculated for each transcription factor in the comparisons described above using the following formula:

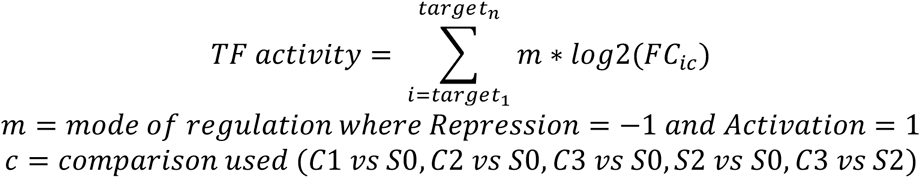

This analysis was performed using R 3.6.2; and heatmaps were generated with the ComplexHeatmap package.

### Immunoblotting

Nuclei and cytosol were prepared from snap frozen tumors using a Thermo nuclear isolation kit according to the supplier’s instructions. Aliquots containing 30 μg (nuclei) or 50 μg protein (cytosol) (assayed by the bicinchoninic acid method) were separated on SDS-polyacrylamide gels containing 5-20% (wt/vol) acrylamide, transferred to nitrocellulose and probed with antibodies as previously described (61).

### Immunohistochemistry, quantitative reverse transcriptase-polymerase chain reaction (qRT-PCR) and DNA methylation analysis

Immunohistochemistry (62), qRT-PCR (63), and assays for DNA methylation (64–66) were performed as previously reported and are described in the Supplemental Methods.

### Clinical Samples

ARIEL2 Part 1 was a multi-center international phase II clinical trial of single-agent rucaparib in patients with platinum-sensitive relapsed ovarian cancer (15). The study was conducted in accordance with the Declaration of Helsinki and Good Clinical Practice Guidelines of the International Conference on Harmonisation. Patients provided written informed consent before participation. Of 196 patients enrolled, four had documented *RAD51C* promoter methylation at some point during the course of their disease. Pretreatment samples from three of these patients were adequate for analysis of *RAD51C* methylation by methylation-sensitive high resolution melting (MS-HRM) analysis (66).

## RESULTS

### Generation of PARPi resistance *in vivo*

To systematically study genomic changes that occur during selection for PARPi resistance *in vivo*, mice bearing PH039, a HGSOC PDX that is extremely sensitive to PARPi but lacks demonstrable mutations in DNA repair genes (67), were treated with multiple cycles of niraparib (**Fig. 1A**). At each step, tumors that regrew were transplanted into additional mice and selected for further resistance.

With the first cycle of PARPi exposure, PH039 shrank dramatically (**Figs. 1B and 1C**) and remained below baseline for >50 days after the end of the treatment (**Fig. 1B**). With the second cycle, shrinkage was again observed, but the tumor regrew more quickly. By the third and fourth cycles, the xenograft grew through niraparib treatment. Histological examination indicated that the tumor recurring after treatment was an adenocarcinoma that was indistinguishable from the source tumor (**Fig. S2A**). Moreover, the signature *TP53* mutation in this tumorgraft model (67) was conserved throughout (**Fig. S2B**). Gene expression profiling (see below) also confirmed that the resistant tumor was of human origin.

### Transcriptome changes associated with mouse passaging and with niraparib resistance

RNA sequencing (RNAseq) was performed on the original passage of PH039 (denoted S0), PH039 passaged without drug treatment (denoted S2 to indicate two passages in untreated mice) and PH039 selected for niraparib resistance (denoted C1-C3 to indicate the number of prior cycles of therapy). The strategy for distinguishing mouse (stroma) vs. human (tumor cell) contributions to levels of each transcript is described in the **Supplemental Results**, **Supplemental Tables S1-S4** and **Supplemental Figs. S1 and S3-S6**. This gene-by-gene analysis not only demonstrated the need to identify and address the nonuniform homologies between human and mouse genes even for PDX tissue with very low mouse tissue infiltration, but also showed that the approach used provided a robust pipeline for quality control in identifying genes disproportionately affected by mouse tissue contamination.

Further analysis demonstrated that certain transcripts changed with repeated passaging in the absence of drug treatment, possibly reflecting adaptation of the tumor to the mouse microenvironment and resulting in diverging and converging patterns of evolution (**Supplemental Results and Figs. S7 and S8**). In addition to these transcript level changes, diverging and converging patterns of upstream pathway regulation were seen when comparing untreated tumor evolution with treated tumor evolution (**Fig. 2A**).

**Figure 2.**
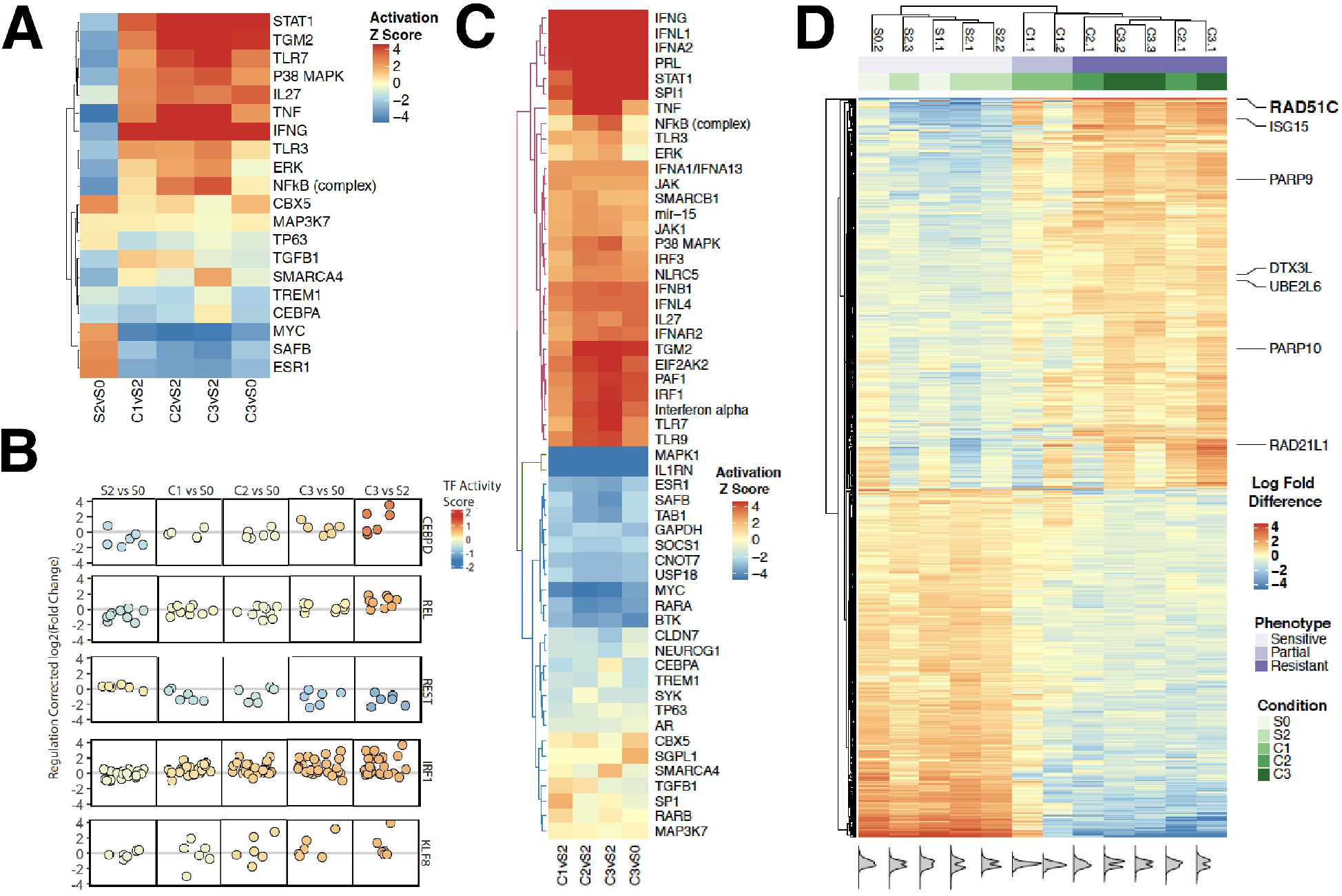
Change in gene expression during PARPi resistance development in PH039. **A,** Evolution of PH039 upstream signaling pathways through selective passaging and drug treatment. Differential expression results (absolute value of log2 fold change >1 and p-value < 0.01) for the pairwise comparisons of samples listed above the heatmap were analyzed through the use of IPA (QIAGEN Inc., https://www.qiagenbioinformatics.com/products/ingenuitypathway-analysis) (44). Upstream regulators changed and detected in all five conditions are shown in the heat-map and colored by z-score. Clustering (rows) represents unsupervised UPGMA clustering on the displayed upstream regulators. **B,** Changes in transcription factor activity scores of the top 5 differentially activated transcription factors between S2 and C3. Each circle represents a target gene that was analyzed. The Y values for each target gene indicate the change in activity, respectively for in the indicated comparison. The colors correspond to the average change in transcription factor activity calculated as described in the Methods. **C,** differential expression results (absolute value of log2 fold change >1 and p-value < .01) for the pairwise comparisons (bottom) were analyzed through the use of IPA (QIAGEN Inc., https://www.qiagenbioinformatics.com/products/ingenuitypathway-analysis (44) and changes in upstream regulators detected in every condition are shown in the heatmap. Color represents the z-score for each upstream regulator in each condition. **D,** Genes differentially expressed between S2 and C3 (absolute value of log2 fold change >1 and FDR < .01) are shown as mean-subtracted log counts per million. Kernel density plots (bottom) are represented for the differentially expressed genes. Hierarchical clustering (columns) was performed using UPGMA on all 13,900 genes passing the filtering threshold. Hierarchical clustering (rows) was performed using UPGMA on the differentially expressed genes shown. Genes with GO terms matching DNA repair are indicated.

To determine whether specific transcription factor activities might contribute to these changes, the TRRUST v2 database was used to examine combined target expression as a measure of putative transcription factor activity for those transcription factors that had at least five identified target genes (**Figs. 2B and S9**). This analysis indicated higher activity scores for CEBPD, which had several target genes upregulated in the resistant PH039 samples, and lower activity scores for REST, which had several target genes downregulated (**Fig. 2B**). In addition, the interferon regulatory factor (IRF) family members IRF9 and IRF1 had higher activity scores in the resistant models when compared to both the parental cell line (S0) and passaged control (S2) (**Figs. 2B and S9B**).

Predicted upstream regulators of the transcriptional changes seen in the resistant tumors were also analyzed by Ingenuity Pathway Analysis (**Fig. 2C**) and showed activation of multiple inflammation-related pathways, including INFG, INFL1, INFA2, TNF, INB1, INFL4, TLR7 and TLR9, as well as downregulation of the SOCS1 pathway, which antagonizes these inflammatory pathways. Interestingly, this activation of inflammatory signaling was identified in two separate analyses, was demonstrated after controlling for mouse tissue contamination in samples that were >50 days from the last PARPi treatment (C1), and was observed in the absence of a functional immune response.

Unsupervised clustering revealed that the untreated tumor tissue, regardless of passage number, clustered with the source PDX (**Fig. 2D**). Additionally, the PDXs that regrew after drug treatment also clustered together, with samples harvested at regrowth after Cycle 1 showing an intermediate gene expression pattern (**Fig. 2D**, columns C1) relative to tumors that regrew after two and three cycles of treatment, which showed a more distinct RNA expression pattern (**Fig. 2D**). Of the greater than 700 differentially expressed transcripts identified in pairwise comparison of S2 and C3, 7 genes (*RAD51C, ISG15, PARP9, DTX3L, UBE2L6, PARP10,* and *RAD21L1*) were associated by gene ontology with DNA repair and are highlighted in **Fig. 2D**. Among these, the top differentially expressed gene was *RAD51C* (log_2_ fold change 8.5),.

### Altered methylation and re-expression of *RAD51C* associates with PARPi resistance

Consistent with the RNAseq analysis, qRT-PCR failed to detect *RAD51C* mRNA in untreated tumor at passage one (S0) and passage six (S5) but revealed readily detectable *RAD51C* message in the PDX that regrew after Cycle 1 (C1) of niraparib, which increased further after additional treatment (C2-C4, **Fig. 3A**). This expression of *RAD51C* mRNA was accompanied by readily detectable RAD51C protein (**Fig. 3B**), which was localized to the nuclei of tumor cells (**Fig. 3C**). In accord with these results, niraparib induced formation of phospho-H2AX foci (**Fig. 3D**), which are consistent with PARPi-induced DNA damage (68), and increased RAD51 foci (**Fig. 3E**), which suggest restoration of HR.

**Figure 3.**
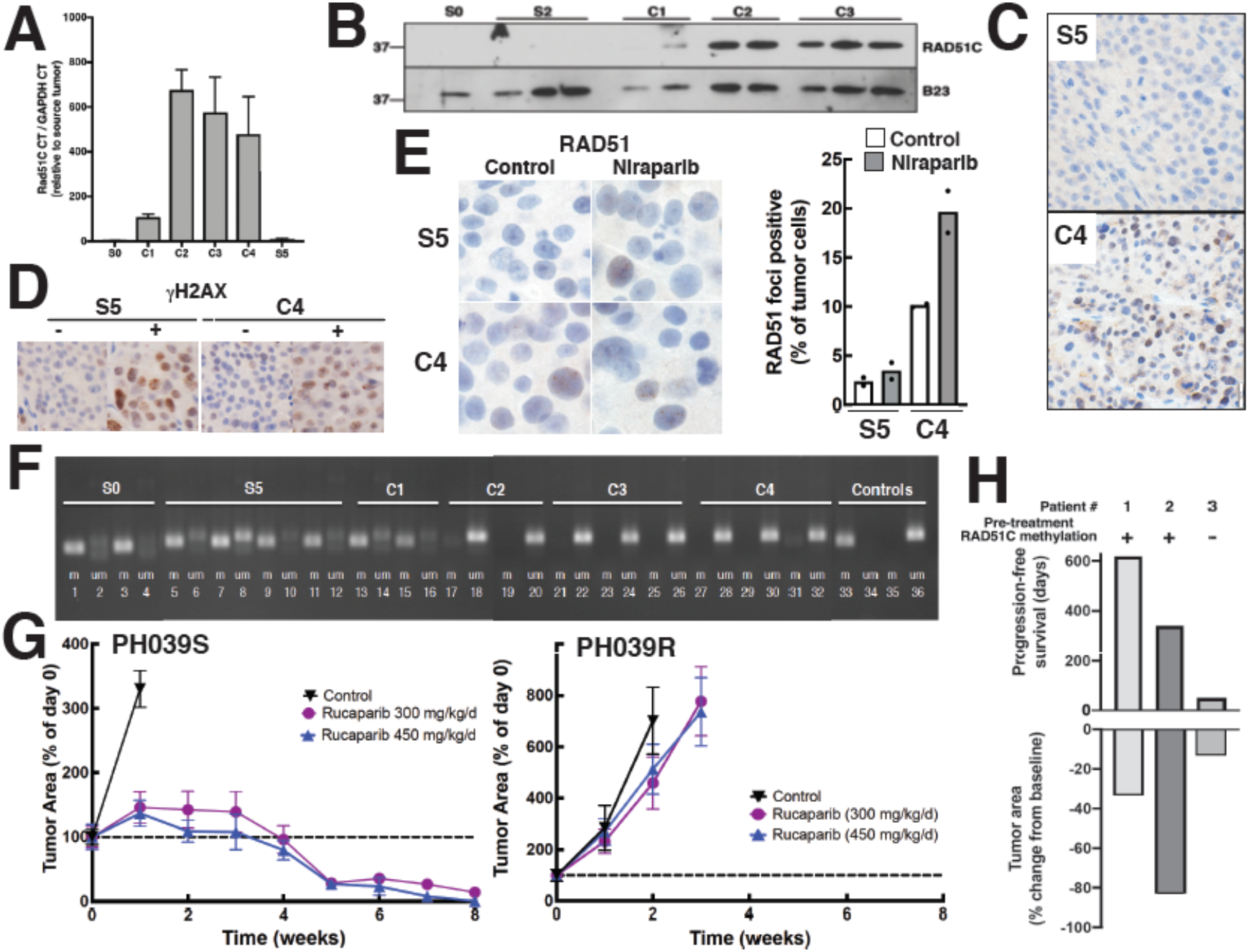
Association between loss of RAD51C methylation and PARPi response. **A,** qRT-PCR results for RAD51C mRNA normalized to GAPDH mRNA. **B,** immunoblot of RAD51C protein across the samples. B23/nucleophosmin served as a loading control. **C,** immunohistochemical RAD51C staining of parental tumor after passaging (S5) versus resistant tumor (C4). **D, E,** phospho-Ser^139^-histone H2AX (D) and RAD51C (E) staining of PH039 (S3) or niraparib-selected PH039R (C4) untreated (-) or after 8 days of niraparib (+). **F,***RAD51C* promoter methylation assessed by bisulfite modification followed by methylation-sensitive PCR in individual PDX samples. Controls were methylated (m) and unmethylated (um) DNA *RAD51C* promoter sequence. **G,** growth of parental (left, S3) or niraparib-selected PH039 (right, C4) without treatment or during treatment with the PARPi rucaparib at 300 or 450 mg/kg/d for 56 days. **H,** three ovarian carcinomas with previous *RAD51C* methylation and platinum-free intervals of 10.8, 32.7 and 24.2 months (Patients 1-3, respectively) were treated with rucaparib monotherapy on the ARIEL2 Part 1 clinical trial after screening biopsies were obtained. HRM analysis on those screening biopsies (**Fig. S10C**, summarized above bars) was compared to days on rucaparib treatment (top bars) and relative change in tumor area as assessed by CT scan using RESIST 1.1 guidelines (bottom bars).

As previously reported, untreated PH039 showed greater than 80% methylation of the *RAD51C* promoter (**Fig. 3F, lanes 1-4**) that persisted during serial passaging (**Fig. 3F, lanes 5-12**). Methylation-sensitive high resolution melting (MS-HRM) analysis demonstrated that this reflected heterogenous methylation of the promoter (**Fig. S10A**). At the end of Cycle 1, the methylation was not detectably different (**Fig. 3F, lanes 13-16; Fig. S10A**). On the other hand, at the end of Cycle 2 when the regrowing PDX was no longer PARPi responsive (**Fig. 1C**), the locus was 95% unmethylated (**Fig. 3F, lanes 17-20**) and was maintained in this unmethylated state through Cycles 3 and 4 of treatment (**Fig. 3F, lanes 21-32; Fig. S10A**). This was further confirmed through allele-specific reduced representation bisulfite sequencing (**Fig. S10B and S10C**), which showed that the diluent-treated PDX contained multiple distinct patterns of *RAD51C* promoter methylation, virtually all of which were lost in the niraparib-selected PDX. Additional analysis of DNA methylation on Illumina EPIC arrays not only confirmed that the *RAD51C* promoter became unmethylated during the course of selection (**Fig. S11A and S11B**), but also revealed that the change in methylation was greater for one of the *RAD51C* promoter probes than for any other probe (**Fig. S11A**). Moreover, changes in methylation were extensive, with over 1100 probes showing a 4-fold change in methylation between S2 and C3 (**Fig. S11A, S11C and S11D**). Among the 700+ genes with altered expression during the course of resistance development (**Fig. 2D**), 64 had a probe with a ≥4-fold change in methylation during resistance development (**Fig. S11C**).

Further xenograft studies revealed that the niraparib-selected PH039 PDX was cross-resistant to the PARPi rucaparib (**Fig. 3G**). Based on these results, we examined the response of ovarian cancers with *RAD51C* promotor methylation identified in ARIEL2 Part 1, a phase II study of the PARPi rucaparib in patients with platinum-sensitive relapsed ovarian cancer (15). Of four cases with *RAD51C* methylation, three had biopsies with sufficient material for MS-HRM analysis harvested immediately prior to rucaparib. Of these three, two cancers showed retention of *RAD51C* methylation at the time of study entry (**Fig. S10D**, patients 1 and 2) and had 33-83% decreases in target lesions by CT on rucaparib treatment, which lasted 339-618 days before progression (**Fig. 3H**). In contrast, the third ovarian cancer had lost *RAD51C* promoter methylation by the time of study entry and had limited shrinkage in response to rucaparib when removed from the study on day 50.

### Dissociation of PARPi and platinum response in PH039

Additional PDX studies demonstrated that PH039R, despite being resistant to niraparib and rucaparib, was still sensitive to carboplatin (**Fig. 4A**). This is in marked contrast to the *BRCA2*-mutated HGSOC PDX PH077, a second PDX model that was also selected by multiple cycles of niraparib treatment. When PH077 became resistant to niraparib (**Fig. 4B,** blue squares), it harbored a secondary mutation that restored the *BRCA2* open reading frame (**Fig. S12A**) and was carboplatin resistant (**Fig. 4B,** green triangles). Likewise, in tissue culture we observed that PARPi and platinum sensitivity varied in parallel when *RAD51C* status was altered (**Fig. 4C and S12B**), in agreement with previous studies (25).

**Figure 4.**
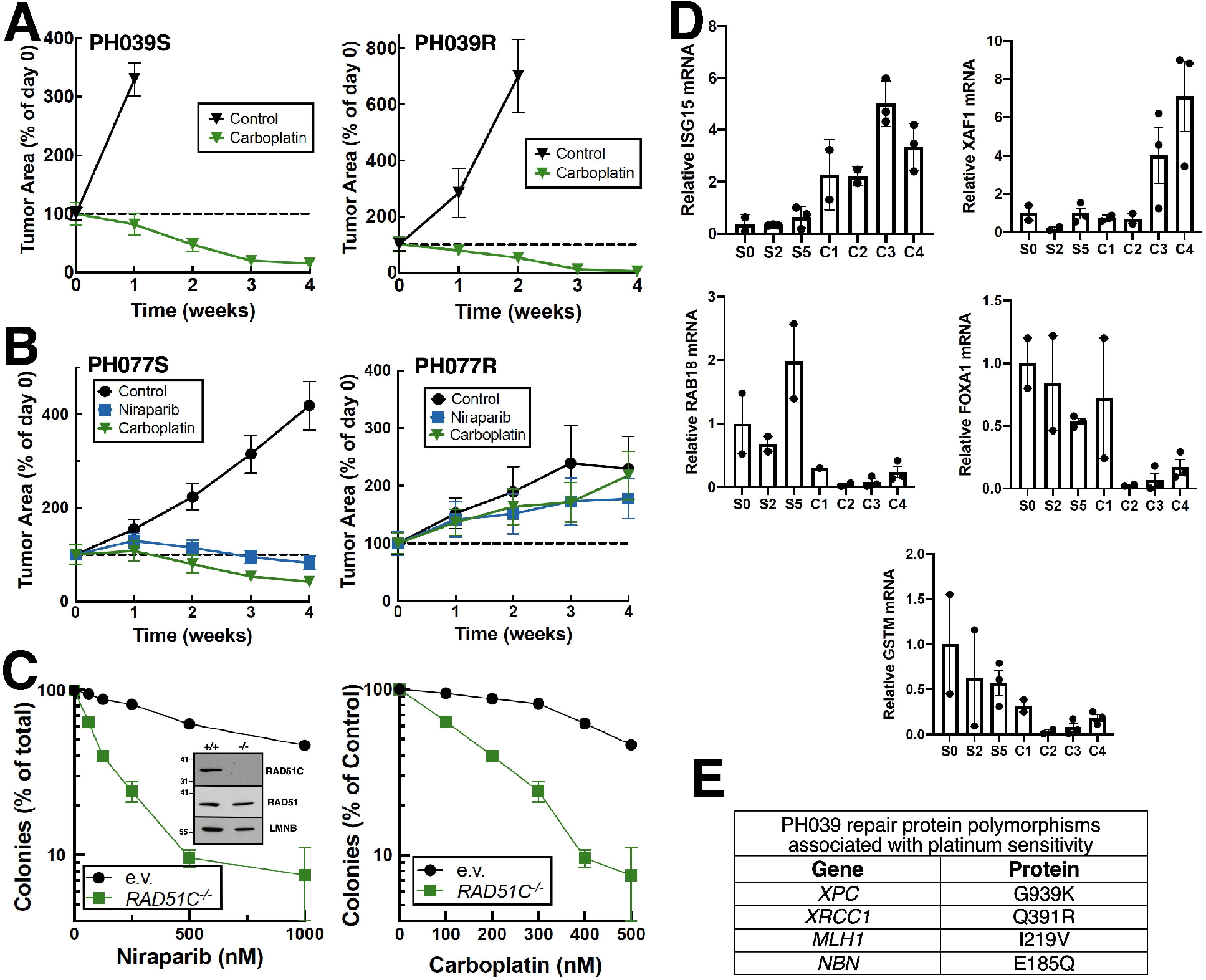
Impact of niraparib selection on platinum sensitivity. **A,** response of PH039S (left) and PH039R (right) to carboplatin 51 mg/kg/week for 4 weeks. **B,** response of PH077S (left) and PH077R (right) to 100 mg/kg niraparib for 28 days or 51 mg/kg/week carboplatin for 4 weeks. **C,** response of parental and *RAD51C^-/-^* Ovcar8 to niraparib and carboplatin in colony forming assays. **D,** qRT-PCR for selected transcripts that have been associated with cisplatin or carboplatin sensitivity. **E,** homozygous genetic polymorphisms observed in PH039 that are associated with high platinum sensitivity.

In order to better understand the unexpected platinum sensitivity of PH039R, we examined DNA sequencing and RNAseq data for mutations and gene expression changes that might be associated with selective platinum sensitivity. Targeted capture and massively parallel sequencing (BROCA analysis, **Table S5**) failed to identify mutations in any of the HR/ovarian cancer susceptibility genes sequenced. On the other hand, further examination of the RNAseq results revealed several changes that might contribute to platinum sensitivity (**Figs. 2D, 4D and S12C**). Among the top 100 differentially expressed genes between PH039S and PH039R, 21 were previously associated with platinum response; and 18 of the changes in expression were reported to convey platinum sensitivity (**Fig. S12C**). In contrast, none of these changes occurred with repeated passaging alone. These changes included upregulation of mRNA encoding the ubiquitin-like protein ISG15 and the XIAP neutralizer XAF1 as well as downregulation of mRNA encoding the glutathione transferase isoform GSTμ, the small G protein RAB18 and the forkhead transcription factor FOXA1 (**Fig. 4D**), all of which have been previously associated with increased platinum sensitivity (69–76). In addition, multiple SNVs that have previously been associated with increased sensitivity to cisplatin or other DNA damaging agents in the clinical setting (77–81) were present at >90% variant allele frequency in the earliest passage PH039 (S0) and persisted in the niraparib-selected PDX (**Fig. 4E**). These included SNVs known to affect proteins in the nucleotide excision repair (XPC G939K), base excision repair (XRCC1 Q391R), mismatch repair (MLH1 I219V), and double-strand break repair (NBN E185Q) pathways. Importantly, each of these variant alleles was confirmed in >90% of RNA reads.

## DISCUSSION

Results of the present study demonstrated that selection for PARP inhibitor resistance in a *RAD51C* promoter-methylated ovarian cancer PDX model is associated loss of *RAD51C* methylation and with development of resistance to multiple PARPis *in vivo*. Similar results were observed in clinical samples from ovarian cancer patients treated on ARIEL2 Part 1, a single-agent phase II clinical trial of the PARPi rucaparib. Interestingly, in the PH039 PDX characterized here, development of PARPi resistance was not accompanied by carboplatin resistance. These observations have potentially important implications for improved understanding of the relationship between PARPi and platinum resistance.

In the present study, we have examined changes in the transcriptome and methylome over the time course of PARPi resistance development in a rare but informative genomic subtype of high grade serous ovarian cancer. Use of a PDX model is ideal for this study, as this number of sequential samples cannot be acquired during the course of clinical treatment. The RNAseq data obtained over the course of resistance development provide a unique look into tumor evolution through sequential passaging with and without drug. Importantly, there are a number of changes during serial passaging even without drug (**Figs. 2D, S7 and S8**), possibly reflecting evolution of this *TP53*-mutant ovarian cancer as it adapts to growth in mice. These observations highlight the potential importance of including both the PDX passaged without drug (S2) and the starting PDX (S0) as controls when examining changes that accompany drug selection.

Comparing the transcriptome of niraparib-selected PH039 to the untreated PDX passaged an equal number of times, the most prominent transcriptional pathway changes reflect increased inflammatory signaling (**Fig. 2A and 2B**). These results are consistent with previous reports that acute PARPi treatment, which is thought to cause release of DNA fragments into the cytoplasm, activates interferon-induced signaling through the cGAS/STING pathway in xenografts and cell culture systems (82–84). Interestingly, however, we observed activation of this signaling over 50 days after PARPi withdrawal (C1 samples, **Figs. 1C and 2**), suggesting that the DNA damage persists long after drug treatment or that PARPis also activate inflammatory signaling through some other persistent process.

The PDX on which these studies were performed, PH039, is among the most PARPi sensitive ovarian cancer PDXs we have examined (33,65). In the therapy naïve model, the *RAD51C* promoter is extensively methylated, a modification that is known to disrupt HR and contribute to PARPi sensitivity (2). Our further analysis demonstrated that this *RAD51C* methylation displays extensive heterogeneity of CpG methylation patterns when analyzed in an allele-by-allele basis (**Fig. S10B**). Upon repeated treatment with niraparib, we observed loss of *RAD51C* promoter methylation accompanied by reappearance of RAD51C mRNA and protein (**Figs. 3A-C, 3F and S10**). Even though the untreated PDX did not contain completely unmethylated alleles (**Fig. S10B**) or a discernible subpopulation of *RAD51C* expressing cells (**Fig. 3C**), partial re-expression of RAD51C and resistance was evident when the tumor grew after the first cycle of chemotherapy. These observations raise the possibility that the diminished *RAD51C* methylation is potentially a dynamic response to PARPi therapy rather than a result of selection of a pre-existing *RAD51C* unmethylated subpopulation of cells.

The change in RAD51C expression was accompanied by changes in expression of over 700 other transcripts. Because over 90% of these changes occurred without alterations in methylation of the corresponding gene promoter (**Fig. S11C**), we examined changes in transcription factor networks that could account for the altered gene expression. As indicated in **Figs. 2B** and **S9**, multiple transcription factors– particularly members of the IRF family– are activated in PARPi resistant PH039.

PARPi sensitivity and platinum sensitivity are generally thought to parallel each other. In PH077, the second ovarian cancer PDX characterized in the present study, this was clearly the case (**Fig. 4B**). This PDX contained a *BRCA2* frameshift mutation at the outset; and selection for niraparib resistance was accompanied by a secondary mutation that restored the *BRCA2* reading frame (**Fig. S12A**) and carboplatin resistance (**Fig. 4B**). In contrast, PH039R was resistant to two different PARPis (**Figs. 1C and 3G**) but sensitive to carboplatin (**Fig. 4A**). Because changes in RAD51C expression clearly affect both PARPi and platinum sensitivity (**Fig. 4C and S12B**), the carboplatin sensitivity of PH039R was unexpected.

Further examination of RNA and DNA sequencing results revealed multiple factors that might, in aggregate, contribute to the platinum sensitivity of PH039R. Among the changes in expression between PH039S and PH039R are many that have been reported to convey platinum sensitivity (**Fig. S12C**), including upregulation of mRNA encoding XAF1 as well as downregulation of FOXA1, RAB18, GSTM, and ISG15, all of which have been confirmed by qRT-PCR (**Fig. 4D**). Among these differentially expressed genes, ISG15 is particularly interesting because its increased expression is i) tightly linked to the increased inflammatory signaling seen after PARPi treatment (**Fig. 2A-2C**) and ii) tied to increased replication stress (85), which would enhance platinum sensitivity. In contrast, the RNAseq analysis did not identify upregulation of any of the ATP binding cassette transporters previously implicated in PARPi resistance (data not shown); and the observation that phospho-H2AX foci form when the resistant PDXs are treated with niraparib (**Fig. 3D**) shows that niraparib still induces DNA damage, further arguing against transport-mediated resistance. On the other hand, previous studies have also shown that cells with nucleotide excision repair (NER) defects are particularly sensitive to cisplatin, which forms the same DNA lesions as carboplatin (86), and resistant to PARPis (31). Examination of PH039R failed to reveal any new mutations in DNA repair genes, including NER genes, compared to the parental PDX. Instead, we identified several SNVs in repair genes that have previously been associated with cisplatin sensitivity in the clinical setting (**Fig. 4E**). Especially noteworthy in this regard is the homozygous XPC G939K alteration, in which the homogeneous A allele seen in PH039 is associated with a higher response rate of non-small cell lung cancer to cisplatin-containing chemotherapy (77). Accordingly, it is possible that the persistent platinum sensitivity of the PH039R PDX reflects the combined effects of several sensitizing features, including i) gene expression changes that counterbalance RAD51C re-expression by increasing platinum sensitivity and ii) the confluence of one or more single nucleotide polymorphisms that also convey platinum sensitivity.

Several observations have previously suggested that the relationship between platinum resistance and PARPi resistance in the clinic might also be more nuanced than generally recognized. First, a large randomized study demonstrated that 67% of suboptimally debulked ovarian cancers exhibit objective responses to cisplatin (87), yet the fraction of ovarian cancers with well-established HR defects is only 40-50% (1,2). These observations raise the possibility that a subset of HR proficient ovarian cancers is platinum sensitive. Second, while the correlation between platinum-free interval (a measure of platinum sensitivity) and % tumor shrinkage on PARPi (a measure of PARPi sensitivity) is statistically significant (30), the correlation is somewhat modest (R^2^ = 0.26), again suggesting that factors beyond HR pathway integrity also might play a role platinum sensitivity. Finally, it has been reported that 40% of ovarian cancers progressing on the PARPi olaparib still respond to subsequent platinum therapy (32), again suggesting that PARPi sensitivity and platinum sensitivity might be dissociated under certain circumstances.

Here we have shown that changes accompanying the development of PARPi resistance in an ovarian cancer PDX with initial *RAD51*C silencing can potentially contribute to retained platinum sensitivity *in vivo*. Whether similar events occur in other ovarian or breast PDXs with *RAD51C* methylation remains to be further investigated. As the present study illustrates, the ability to examine sister tumors for sensitivity to a variety of agents represents one of the clear-cut benefits of PDXs for understanding patterns of cross resistance.

## Supporting information

Supplemental methods, results, tables and figures

## ACKNOWLEDGEMENTS

We thank David Toft for his kind gift of H90-10 antibody and Alan D’Andrea for helpful discussions. This work was supported in part by grants from the NIH (P50 CA136393 to S.H.K., L.M.K., H.L., and S.J.W.; F30 CA213737 to C.D.M.; T32 to GM072474 to R.M.H. and R.L.K.; T32 GM085641 to T.M.W.), Ovarian Cancer Research Alliance (E.M.S., P.H. and S.H.K.), Stand Up to Cancer (A.E.W.H., E.M.S., S.J.W. and S.H.K.), the Stafford Fox Medical Research Foundation (CLS, OK, MW, KN), and the National Breast Cancer Foundation of Australia (AD) as well as fellowship support from the Mayo Foundation for Education and Research (R.M.H., C.D.M., T.M.W., R.M.K., and A.V.).

## AUTHOR CONTRIBUTIONS

**Conceived and designed project:** R.M.H., C.D.M., P.H., S.H.K.

**Conducted experiments:** R.M.H., K.N., R.L.K., A.V., X.H., N.M.P., X.W.M., M.R.R., P.A.S., K.S.F., K.L.P., A.D., J.M.W., S.H.K.

**Analyzed data:** R.M.H., C.D.M., K.N., C.C., T.M.W., O.K., C.A.R., M.W., H.L.

**Provided critical reagents:** K.K.L., T.C.H., I.A.M.

**Project supervision:** M.W., C.L.S., A.E.W.H., L.M.K., E.M.S., H.L., S.J.W. and S.H.K.

**Initial manuscript preparation:** R.M.H., C.D.M. and S.H.K.

**Manuscript editing:** All

