## Supplemental methods, results, tables and figures for "Characterization of a *RAD51C*-Silenced High Grade Serous Ovarian Cancer Model During PARP Inhibitor Resistance Development"

#### *In vivo* efficacy of PARP inhibitors and chemotherapy

For all PDX studies, cryogenically preserved human ovarian cancer tumors were rapidly thawed and reestablished in female SCID Beige mice (C.B-17/IcrHsd-*Prkdc*$^{scid}$*Lyst*$^{bg-}$J; Envigo, Indianapolis, IN) as previously described (1). Briefly, 0.1-0.2 cc of minced tumor was prepared in 1:1 ratio with McCoy's 5A Modified Medium(Cat # MT-10-050-CV, Corning Life Science) before intraperitoneal injection. Control tumors (PH039S and PH077S) and resistant tumors (PH039R and PH077R) were matched for passage number. When tumor cross-sectional areas reached 0.3–0.5 $cm^2$ by transabdominal ultrasound, mice were randomized to treatment arms. The rucaparib studies were conducted at 300 or 450 mg/kg by daily gavage in 0.1% methylcellulose for 56 days with 5 mice per group. Control mice (n=5) served as a reference for growth kinetics, but sensitivity to treatment was defined as regression below baseline. Ultrasound measurements were taken weekly and plotted as the mean tumor area percent relative to the starting baseline size. Saline or carboplatin at 51 mg/kg, a dose based on standard platinum-doublet dosing (1) but with paclitaxel was omitted to permit assessment of sensitivity to platinum alone, were given to 10 mice/group by weekly IP injection for four weeks, followed by a week of observation before the study endpoint. Niraparib treatments in 6-7 mice/group were administered at 100 mg/kg by daily gavage for 28 days and tumor size was assessed as above for the carboplatin treatments.

#### BROCA Analysis

DNA was harvested from PDX models using an All Prep DNA/RNA MiniKit (Qiagen) and assayed for mutations in genes involved in DNA repair (**Table S5**) by massively parallel DNA sequencing as previously described (2). Mutations were considered deleterious if they were truncating or were missense mutations with evidence of functional compromise. Sanger sequencing was used to confirm deleterious mutations.

#### DNA methylation

250 ng of DNA was bisulfite converted (EZ Methylation Direct kit, Zymo Research, Irvine, CA) and evaluated with methylation sensitive PCR for *RAD51C* as previously described (3,4). In brief, PCR was performed with primers for the methylated (M) reaction (sense-5′-TGtAAGGttCGGAGtttCGTGC-3′ and antisense- 5′-TCGCTaaaaCGTaCGaCGTaACG-3′, 85 nt) and unmethylated (UM) reaction (sense-5′-GtGtAA-AGtTGtAAGGtttGGAGttttGTGtG-3′ and antisense-5′-CaCACaCCCTCaCTaaaaCaTaCaaCa-TaACa-3′, 103 nt). Methylated and unmethylated DNA (#D5014) was used as positive M control (M ctrl) and UM control (UM ctrl) for bisulfite conversion and validation of amplicon size. Water ($H_2O$) was substituted for DNA to rule out cross-contamination of samples. *RAD51C* methylation was confirmed by MS-HRM, as previously described (5).

### EPIC array analysis

DNA methylation assays were performed as per the standard manufacturer’s protocol (http://www.illumina.com/). In brief, 500 ng of DNA extracted from the PDX and murine sample were treated with sodium bisulfite to convert cytosine to uracil. The 5-methyl cytosine remains unreactive to sodium bisulfite. The DNA is then hybridized to the EPIC BeadChip. After washing off unhybridized DNA, a single base extension was recorded to calculate the methylation level at the CpG probe site. DNA methylation assays were performed at The Walter and Eliza Hall Institute of Medical Research (Australia). Raw intensity data files (idat files) for both PDX and murine samples were normalized using the within-array normalization method SWAN (subset-quantile within array normalization) function in the missMethyl R package (6-8). The Limma package in R was then used to perform differential methylation analysis comparing treated and untreated

PH039 samples using a log fold change threshold > 2 (9). EPIC array probes that overlap human-mouse were excluded (10). Pathway enrichment analysis was performed using the ReactomePA, and data were plotted using ggplot2 and complex heatmap R packages (http://www.bioconductor.org/packages/ReactomePA).

#### Whole Exome Sequencing

Paired-end libraries were prepared using 1.0 µg of genomic DNA with the Agilent Bravo liquid handler according to instructions from Agilent Technologies. The concentration and size distribution of the completed libraries was determined using an Agilent Bioanalyzer DNA 1000 chip (Santa Clara, CA) and Qubit fluorometry (Invitrogen, Carlsbad, CA). Whole exome capture was carried out using 750 ng of the prepped library following the protocol for the SureSelect Human All Exon v5 + UTRs 75 MB kit (Agilent). The purified capture products are then amplified using the SureSelect Post-Capture Indexing forward and Index PCR reverse primers (Agilent) for 12 cycles. Libraries were sequenced at an average coverage of 80 x following Illumina's standard protocol using the Illumina cBot and cBot Paired end cluster kit version 3. The flow cells were sequenced as 101 x 2 paired-end reads on an Illumina HiSeq 2000 using TruSeq SBS sequencing kit version 3 and HCS v2.0.12data collection software. Base-calling was performed using Illumina's RTA version 1.17.21.3 (11).

FASTQ alignment was conducted with bwa-mem v0.7.12 against GRCh38.86 human reference genome build using default parameters and realignment was performed using GATK (v3.4-46) (12). To avoid confounding of results by mouse reads, variants detected in RNAseq were manually inspected in aligned BAM files.

#### Quantitative reverse transcriptase-polymerase chain reaction (qRT-PCR)

RNA was isolated from snap-frozen tissue using in RNeasy Plus Minikit (Qiagen). qRT-PCR was performed in triplicate using 100 ng RNA and TaqMan RNA-to-$C_T$ 1-Step Kit (Applied Biosystems, Carlsbad, CA) per the supplier's instructions. Using probesets for GAPDH (4352665) from Life Technologies) and RAD51C (Hs04194939_s1), IGS15 (Hs01921425_s1), RAB18 (Hs04981645_s1), FOXA1 (Hs04187555_m1) and GSTM1 (Hs01683722_gH) from Applied Biosystems (Carlsbad, CA), PCR was performed on a CFX384 Real Time System (C10000 Touch Thermal Cycler, BioRad, Hercules, CA) using a program consisting of 48°C for 15 min, 95°C for 10 min, then 40 cycles of 95°C for 15 sec and 60°C for 1 min. Data were analyzed using the following equations: $\Delta\Delta C_t = \Delta C_t$(sample)- $\Delta C_t$(endogenous control); and Fold Change = $2^{-\Delta\Delta Ct}$, and assessed for relative change as indicated in the various figures.

#### RAD51C and RAD51 immunohistochemical (IHC) staining

Tissues harvested from mice were fixed overnight in buffered formalin (Fisher Scientific; #23-011-120), processed, and stained with hematoxylin and eosin in the Histology Core Facility at Mayo Clinic (Scottsdale, AZ). PE04 ovarian cancer cells transfected by electroporation with control siRNA or pooled RAD51C siRNAs were formalin fixed and paraffin embedded to serve as positive and negative controls, respectively. Tissue sectioning and IHC staining were performed at the Pathology Research Core (Mayo Clinic, Rochester, MN) using the Leica Bond RX stainer (Leica). For RAD51C staining, FFPE tissues were sectioned at 5 microns, subjected to antigen retrieval for 30 min using Epitope Retrieval 2 (EDTA; Leica), and incubated for 5 min in Protein Block (Dako). The RAD51C primary antibody (Novus NB100-177) diluted 1:100 in Background Reducing Diluent (Dako) was incubated with slides for 30 min.

Detection used a Polymer Refine Detection Kit (Leica, Allendale, NJ). Slides were counterstained on-line with hematoxylin for 5 min followed by several rinses in 1X Bond wash buffer and distilled water. Slides were then removed from the stainer, rinsed in tap water for 5 min, dehydrated in increasing concentrations of ethanol and cleared in 3 changes of xylene prior to permanent coverslipping in xylene-based medium.

RAD51 staining was performed with the following changes: i) Antigen retrieval was performed using Epitope Retrieval 1 (Leica), ii) anti-RAD51 clone EPR4030(3) (Abcam, ab133534) was diluted 1:20,000 in Background Reducing Diluent (Dako), and iii) PE04 and PE01 cells harvested 6 h after 1000 cGy ionizing radiation were formalin fixed and paraffin embedded to serve as positive and negative controls, respectively. Ten random fields from each slide were examined for foci on a Zeiss Axiophot microscope equipped with a 63X Zeiss N.A. 1.4 lens and photographed as Z stacks.

**Tissue culture, gene interruption and colony forming assays**

Ovcar8 cells, provided by Dominic Scudiero (NCI Frederick) and authenticated by short tandem repeat analysis in the Mayo Clinic Cytogenetics Core Facility, were cultured in RPMI 1640 medium containing 10% heat-inactivated fetal bovine serum, 10 μg/ml insulin, 50 U/ml penicillin G, 50 μg/ml streptomycin, and 2 mM glutamine (medium A) at 37 °C in an atmosphere of 5% $CO_2$.

For siRNA experiments, aliquots containing 8 million Ovcar8 cells were transfected with Dharmacon (Lafayette, CO) pooled siRNAs to RAD51C (L-010534-00-0010) or nontargeting control by electroporation (2 μM siRNA in 200 μl volume; two 10 mSec 280 V pulses in a BTX830 square wave electroporator) on two consecutive days. Beginning 48 h after the second electroporation, aliquots were harvested for preparation of nuclear extracts as described in the main text or plated (300 siControl cells or 450 siRAD51C cells in 3 ml medium A per 35 mm well or dish), incubated for 4-16 h to allow cells to adhere, and treated for 48 h with niraparib or cisplatin (added from 1000X stocks in DMSO). Cells were then washed and incubated under drug-free conditions for 8-10 days to allow colony formation.

*RAD51C*$^{-/-}$ Ovcar8 cells were generated by CRISPR/Cas9 technology (13) using a lentiviral construct (Genescript AF029669 gRNA1) targeting nt 929-948. Two days after transduction, cells were selected with 3 μg/ml puromycin. Pooled cells that repopulated the culture were cloned by limiting dilution and assayed for RAD51C expression in nuclei by immunoblotting. Cloned *RAD51C*$^{-/-}$ cells or pooled cells transduced with empty vector were subjected to colony forming assays as described with 300 (empty vector) or 450 cells (*RAD51C*$^{-/-}$) plated and with continuous drug exposure.

### SUPPLEMENTAL RESULTS

#### Considerations of multi-mapping in the analysis of the PDX RNAseq data

Due to the high homology between human and mouse genomes, mapping RNAsequencing data to the appropriate organism (mouse or human) can present a challenge. One commonly used PDX specific software is Xenome (14), which uses a k-mer decomposition to classify reads as human, both, or mouse. Processed reads can then be mapped normally. Conway *et al*. also describe a method to use TopHat (15,16) to differentiate which reads fall to which genomes and which genes are classified as aligning to both.

Two newer approaches have aimed to differentiate the genes that are ambiguous between mouse and human aligners. Disambiguate (17) proposed using read quality alignment to help differentiate reads mapped to both genomes. Another recent approach by Callari *et al.* uses a combined human-mouse reference genome for alignment, cutting the computational cost of Disambiguate with similar performance (18). In their comparison between pipelines, however, Callari *et al.* show that even in RNAseq data that is purely human or purely mouse, multi-mapping continues to occur with all software.

Whether or not multi-mapping will affect the differential expression results for a given PDX study depends on the underlying distribution of the multi-mapping phenomenon. For example, if there are several genes with high homology that are expressed at very low levels in the human tumor but at high levels in the mouse stroma, it is conceivable that these genes could be identified as differentially expressed due to changes in mouse stroma (either characteristics or percent infiltration) rather than changes in the tumor. Similarly, if there is a gene with high homology between mouse and human that is expressed at high levels in the mouse stroma relative to the human tumor, the mouse stroma expression could cover genes that are differentially expressed within the tumor.

To try to sort reads based on species, we took the approach outlined in **Figure S1**. In brief, the raw reads were aligned to either the human or mouse genome. The reads that were aligned to the mouse genome were then removed and the unaligned reads were mapped to the human genome. The percentage of read removal is shown in **Table S1**. Any genes for which the read mapping was changed by two-fold after mouse removal in any sample were flagged as having potential multi-mapping. The upstream method of dual genome alignment with TopHat (15,16) to identify reads mapping to both genomes is analogous to the method outlined in the Xenome paper (14). The read removal results are shown in **Tables S1** and **S2**. On average, 7.39% of reads mapped to the human annotated transcriptome were removed after the reads mapping to the mouse genome were removed. Of the reads that mapped to the annotated mouse transcriptome, an average of 46.3% were removed after reads mapping to the human genome were removed. Taken together, this indicates that of the reads mapped to the mouse genome, almost half display multi-mapping with the human genome,whereas less than 10% of the human alignment multi-maps to the mouse genome.

In order to understand the distribution of the multi-mapping, a comparison of read counts per gene between the human only alignment and human alignment after mouse removal was visualized. As shown in **Figure S3**, several genes with high expression were identified as differentially expressed between the human only pipeline and the pipeline after mouse read removal. Thus, even with low mouse read mapping there were genes that were disproportionately affected by multi-mapping. In order to study this phenomenon so we could adequately control for it, genes that showed significant differences in read count between the mouse and human study were flagged (see **Figure S3** for criteria). A subset of this list for which the average read count was greater than two and a gene was identified in more than two samples is shown in **Table S3**. Some important genes in ovarian cancer were immediately identified, including PTEN (frequently deleted) and ZEB2 (involved in epithelial to mesenchymal transition.) In addition, lesser studied genes such as CAMTA1 were identified, as was RAB18, a transcript identified as differentially expressed between S2 and C3 ($\log_2 FC = -5.6$, fdr <0.0001).

Using the BLAST alignment tool (19), mRNA sequences from these genes as well as for RAD51C and CDH1 (not flagged for multi-mapping) were aligned to the mouse transcriptome. The results are shown in **Table S4**. The BLAST alignments for these genes as well as the raw read counts aligned to each are shown in **Figures S4** (ZEB2 and PTEN), **S5** (CAMTA1 and RAB18), and **S6** (RAD51C and CDH1).

The cause of multi-mapping can be appreciated by careful examination of the phenomenon for each gene. Many more reads from ZEB2 and PTEN map to mouse than to human and the percent identity between human and mouse respectively is 93 and 90 percent (**Table S4**). Furthermore, almost all genes that mapped to human are removed after mouse alignment. This indicates that the most likely source for these reads is actually the mouse stroma. CAMTA1, on the other hand, has the highest percent identity (98.54%) with 95% coverage (**Table S4**). For CAMTA1, it is difficult to tell whether the identity of the reads is more likely human or mouse, as the percent of read removal is so high (**Figure S5**). However, it is clear that CAMTA1 does have about a third of reads uniquely human and almost no reads uniquely mouse. RAB18 (**Figure S5**) has 91.99% identity, however the coverage is only 17%. This gene is identified because in samples C2 and C3, this gene had more than two-fold greater counts in the human sample before mouse read removal than after. The most likely interpretation is that, when the human RAB18 reads drop to extremely low numbers, the mouse RAB18 (which is reasonably highly expressed) begins to become a large percentage of the detected RAB18 reads. Finally, the two genes not flagged as multiply mapped (RAD51C and CDH1) show lower percent identity (88% and 80% respectively) and low coverage (42% and 56% respectively). Direct checking of the mapped reads reveals little change between the pipelines for human and mouse removal (**Figure S6**).

Taken together, these results demonstrate a robust pipeline for quality control in identifying genes that are disproportionately affected by mouse tissue contamination and pipeline differences. Even for PDX tissue with very low mouse tissue infiltration, these results underscore the need for appropriate controls in PDX sequencing studies due to the nonlinear homologies between human and mouse genes.

### Evolution of PH039 Signaling Pathways through Selective Passaging

In order to understand how to best perform the differential expression analysis over the course of resistance development in PH039, the relationship between the source tumor (S0) and the cyclically passaged untreated tumor (S2) was compared with the resistant tumor (C3). **Figure S7** shows genes differentially expressed between S0 and S2 ($|\log_2 FC| > 1$ and p-value < 0.01) with C3 shown for comparison. Black labeled genes indicate genes that were differentially expressed between C3 and S2 with $|\log_2 FC| > 2$ and p-value < 0.01) and red labeled genes indicate genes flagged as differentially expressed between mouse and human pipelines. On visual inspection, many of the genes that were differentially expressed between C3 and S2 and between S0 and S2 may actually represent genes that are unchanged between S0 and C3.

In order to investigate the patterns of gene evolution, a plot of converging and diverging genes is shown in **Figure S8**. Genes that were both significantly increased (or decreased) from S0 to C3 and from S0 to S2 are shown and represent convergent evolution. These genes may reflect the selective pressure of passaging in the mice rather than drug effect. Divergent evolution is captured as the genes that were significantly changed from source in opposite directions. These genes represent genes that were undergoing opposite selective pressures through passaging. Finally, pathway analysis by Ingenuity Pathway Analysis (IPA) of the predicted upstream regulators identified across all comparisons for the genes differentially expressed between each condition set is shown in **Figure 2A** (QIAGEN Inc., https://www.qiagenbioinformatics.com/products/ingenuitypathway-analysis) (20). Using S0 versus C3 as a comparison, pathways can be identified that are significantly changed through drug treatment versus those that are changed through mouse pressure alone. For example, ERK signaling is predicted to be up-regulated between S2 and C3 and down-regulated from S0 to S2. However, in the S0 versus C3 results, there is little predicted change. This likely indicates that the pathway up-regulation seen in S2 versus C3 is an artifact of the divergence of S2 and S0 rather than selective drug pressure. Similarly, enrichment of CBX5 is seen in S0 versus C3 and S0 versus S2, but not in S2 versus C3. This pattern, though different from that of ERK, is also indicative that the CBX5 predicted upstream activation is likely due to selective pressure during passaging in the mouse. STAT1, on the other hand, is predicted to have down-regulated signaling in S0 versus S2 and up-regulated signaling in S2 versus C3. Again, by observing the S0 versus C3 comparison, we see that the pathway down-regulation between S0 and S2 is not significantly contributing to the up-regulation seen between S2 and C3. Therefore, it can be concluded that the predicted STAT1 activation is likely changed as a result of drug effect rather than selective pressure by mouse passaging.

Taken together, these results demonstrate that comparing cyclically passaged untreated tumors against the cyclically passaged drug treated tumors can help identify patterns of gene evolution as well as provide an important control for understanding pathway analysis results.

### Table S1
### Reads Mapped to Human Transcriptome
### Before and After Mouse Alignment Removal

| Sample | Total Reads Mapped to HG38 | Total Reads Mapped to HG38 after Mouse Read Removal | % Read Removal |
|---|---|---|---|
| **S0.1** | 68559822 | 64234213 | 6.31 |
| **S0.2** | 82591769 | 77267716 | 6.45 |
| **C1.1** | 77665230 | 71187348 | 8.34 |
| **C1.2** | 80604229 | 74535260 | 7.53 |
| **C2.1** | 64806333 | 61140036 | 5.66 |
| **C2.2** | 69409509 | 65268406 | 5.97 |
| **C3.1** | 74055199 | 69082150 | 6.72 |
| **C3.4** | 90870530 | 84831623 | 6.65 |
| **C3.3** | 75425752 | 70393032 | 6.67 |
| **S2.1** | 68889027 | 63086892 | 8.42 |
| **S2.2** | 64156503 | 59209277 | 7.71 |
| **S2.3** | 86628922 | 76014479 | 12.25 |

Total number of reads mapped to the UCSC annotated human transcriptome (HG38) before and after read removal from mouse (MM10) alignment. This represents the upper bound of mouse read contamination expected in human transcriptome results. (See Table S2 for analogous mouse read comparison.)

### Table S2
### Reads Mapped to Mouse Transcriptome Before and After Human Alignment Removal

| Sample | Total Reads Mapped to Mouse | Reads Mapped to Mouse after Human Removal | % Read Removal |
|---|---|---|---|
| S0.1 | 7174132 | 2116314 | 29.5% |
| S0.2 | 12718711 | 5401617 | 42.5 % |
| S2.1 | 15218863 | 7405806 | 48.7% |
| S2.2 | 11983372 | 4521993 | 37.7 % |
| S2.3 | 22754938 | 5829572 | 25.6% |
| C1.1 | 23191089 | 11974165 | 51.6% |
| C1.2 | 21782124 | 12228047 | 56.1 % |
| C2.1 | 8610509 | 3643086 | 42.3 % |
| C2.2 | 18460135 | 12680920 | 68.7 % |
| C3.1 | 19687127 | 12554130 | 63.8% |
| C3.2 | 16221681 | 7038493 | 43.4 % |
| C3.3 | 18580929 | 10358086 | 55.7 % |

Total number of reads mapped to the UCSC annotated mouse transcriptome (MM10) before and after read removal from human (HG38) alignment. This represents the upper bound of human read contamination expected in mouse transcriptome results. (See Table S1 for analogous human read comparison.)

### Table S3
### Flagged Genes for Mouse/Human Pipeline Differences

| | | | | | |
|---|---|---|---|---|---|
| CAMTA1 | 12 | DMD | 10 | PRKD1 | 3 |
| RAP1A | 12 | GRIA3 | 10 | GOLGA6L17P | 3 |
| EIF5AL1 | 12 | CSNK2A3 | 8 | CPEB1 | 3 |
| PTEN | 12 | IGF1 | 8 | OSR1 | 3 |
| SPI1 | 12 | NEXN-AS1 | 7 | AHCTF1P1 | 3 |
| RNU2-2P | 12 | NOVA1 | 7 | FN1 | 3 |
| TUBA1A | 12 | ANTXR2 | 7 | BMP2 | 3 |
| RNU4-2 | 12 | EBF1 | 7 | TMEM183B | 3 |
| KPNA3 | 12 | EBF3 | 6 | HOXA7 | 3 |
| RN7SL1 | 12 | HOXC10 | 6 | PMS2P7 | 3 |
| RN7SL2 | 12 | HOXC8 | 6 | HGF | 3 |
| MAF | 12 | DDT | 6 | CAV1 | 3 |
| MIR3064 | 12 | EPB41L2 | 6 | RPS26P11 | 3 |
| ZEB2 | 12 | GTF2IP1 | 6 | FAM45B | 3 |
| RNA18S5 | 12 | ZFHX4 | 6 | ARHGEF6 | 3 |
| RNA28S5 | 12 | ZEB1 | 5 | DDR2 | 2 |
| MEF2C | 12 | HIC1 | 5 | FAM78B | 2 |
| SPARC | 12 | IKZF1 | 5 | PRG4 | 2 |
| RN7SK | 12 | SH2D3C | 5 | PRKG1 | 2 |
| SYNCRIP | 12 | FHL1 | 5 | EDNRB | 2 |
| CELF2 | 11 | RAB18 | 4 | HOXD10 | 2 |
| CSNK1A1L | 11 | RPL13AP5 | 4 | PCBP3 | 2 |
| CDH11 | 11 | HOXC6 | 4 | GATA2 | 2 |
| MIR5047 | 11 | SMG1P2 | 4 | GRIA2 | 2 |
| DAB2 | 11 | UBE2MP1 | 4 | PPP2R2B | 2 |
| PRRX1 | 10 | COL1A1 | 4 | ETV1 | 2 |
| RNU4-1 | 10 | FAP | 4 | ST7-OT3 | 2 |
| BMP4 | 10 | CACNA1D | 4 | RNF5P1 | 2 |
| MAP2K4 | 10 | FST | 4 | RUNX1T1 | 2 |
| TIMP3 | 10 | RNVU1-7 | 3 | HS6ST2 | 2 |
| SFRP1 | 10 | FLI1 | 3 | | |

Genes with $|\log_2(1+Y) - \log_2(1+X)| > 1$ and $\frac{1}{2}(\log_2(1+Y) + \log_2(1+X)) > 2$, where X read count to a particular gene after mouse read subtraction and Y is the read count to a particular gene without mouse read subtraction (see Figure S4). The numbers to the right of the gene indicate the number of samples for which this criterion was met. For example, the genes followed by the number 12 had $\log_2(1+Y) - \log_2(1+X) > 1$ for every sample whereas genes marked by 2 only met this criteria in two of the 12 samples. Four genes that appeared in the differential expression analysis between C3 and S2 are highlighted. Blue indicates abs(lfc)>1(HOXC6, RNA18S5, and SPI1) and red indicates abs(lfc)>2 (RAB18).

**Table S4**
**BLAST Results for Select Genes**

| Human Gene | Human Accession Number | Query Cover-age | Percent Identity | Mouse Accession Number | Mouse Gene |
|---|---|---|---|---|---|
| *CAMTA1* | NM_015215.4 | 95% | 98.54% | NM_001081557.3 | *Camta1* |
| *ZEB2* | NM_014795.4 | 94% | 93.23% | NM_001355289.1<br>NM_001355288.1<br>NM_015753.4 | *Zeb2* |
| *RAB18* | NM_021252.5 | 17 % | 91.99% | NM_181070.6 | *Rab18* |
| *PTEN* | NM_000314.7 | 57% | 90.63% | NM_008960.2 | *Pten* |
| *RAD51C* | NM_058216.3 | 42% | 87.57 % | NM_053269.4 | *Rad51c* |
| *CDH1* | NM_004360.5 | 56% | 79.75% | NM_009864.3 | *Cdh1* |

BLAST results (19) for alignment of the human reference mRNA sequence for each gene in the table downloaded from the NCBI database (21) against the mouse RefSeq RNA database. Query coverage and percent identity are shown with the corresponding best alignment result from the mouse transcriptome. The e-value was 0 for all of the top hits shown. Corresponding figures are found in Figures S4-S6.

### Table S5. Genes Assessed for Mutations in BROCA Analysis

| | | | |
|---|---|---|---|
| ATM | ERCC3 | MLH1 | RAD51D |
| ATR | ERCC4 | MRE11A | RBBP8 |
| BABAM1 | ERCC5 | MSH2 | RECQL |
| BAP1 | ERCC6 | MSH6 | RIF1 |
| BARD1 | EZH2 | NBN | RINT1 |
| BLM | FAM175A | NEIL1 | SLX4 |
| BRCA1 | FANCA | PALB2 | SMARCA4 |
| BRCA2 | FANCB | PARP1 | TOPBP1 |
| BRCC3 | FANCC | PAXIP1 | TP53 |
| BRE | FANCE | PIK3CA | TP53BP1 |
| BRIP1 | FANCF | PMS2 | UBE2T |
| CDH4 | FANCG | POLD1 | UIMC1 |
| CDK12 | FANCI | POLE | USP28 |
| CHD4 | FANCL | POLQ | WRN |
| CHEK1 | FANCM | PPM1D | XPA |
| CHEK2 | GEN1 | PTEN | XPC |
| DCLRE1C | HELQ | RAD50 | XRCC2 |
| DDB1 | ID4 | RAD51 | XRCC3 |
| ERCC1 | LIG4 | RAD51B | XRCC4 |
| ERCC2 | MAD2L2 | RAD51C | XRCC5 |
| | | | XRCC6 |

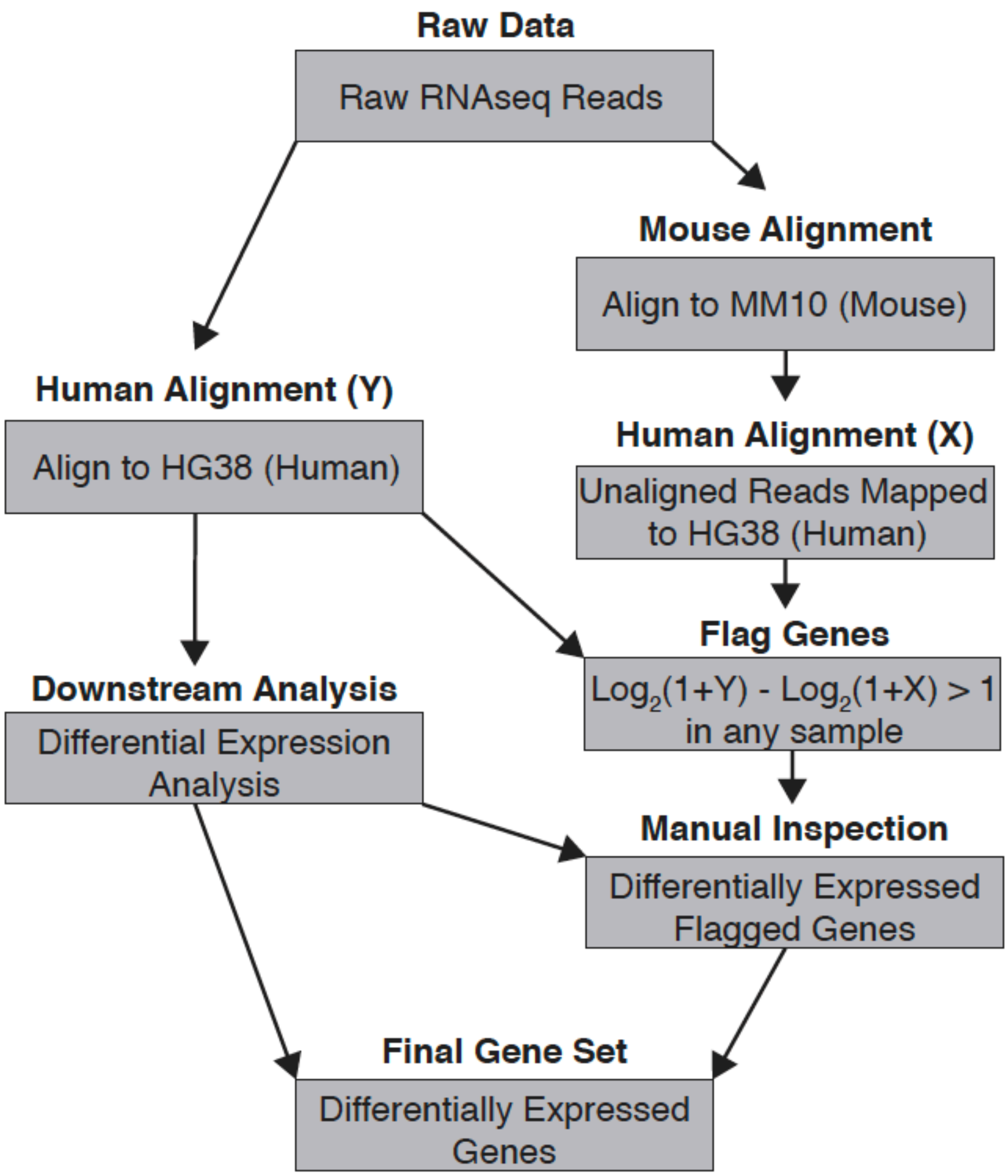


**Figure S1. RNAseq pipeline (related to Fig. 2).** Raw reads were aligned to human HG38 (Y) or mouse MM10 with TopHat. Unaligned reads after TopHat MM10 alignment were converted back to FASTQ format with Picard (http://broadinstitute.github.io/picard). Reads after mouse removal were aligned to HG38 using TopHat. FeatureCounts (22) was used to assign a read count to each gene based on the UCSC gene annotations associated with HG38. Genes where the counts from human only alignment (Y) and the mouse-removal human alignment (X) were evaluated by the formula $\log_2(1 + Y) - \log_2(1 + X)$ and were flagged if this quantity was greater than 1 in any sample. Differential expression analysis proceeded for the human only alignment and all genes identified as both differentially expressed and flagged were individually evaluated.

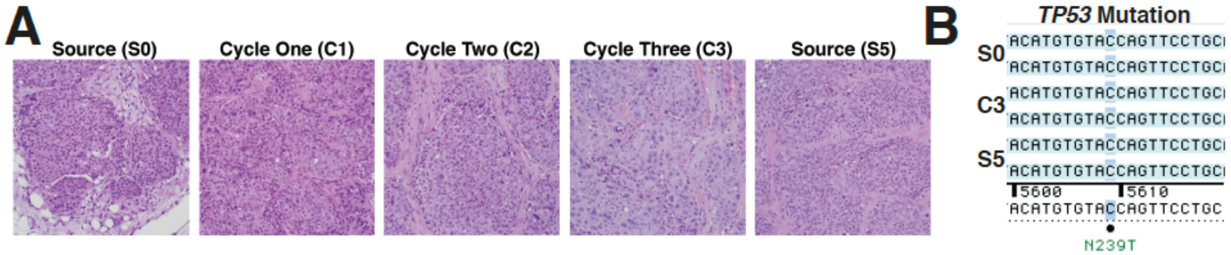


**Figure S2 (related to Fig. 1). Further characterization of niraparib-resistant PH039 xenografts. A,** adenocarcinoma histology was confirmed throughout the development of resistance. **B,** sequencing confirmed the presence of the patient tumor *TP53* mutation throughout both resistance development and source tumor passaging.

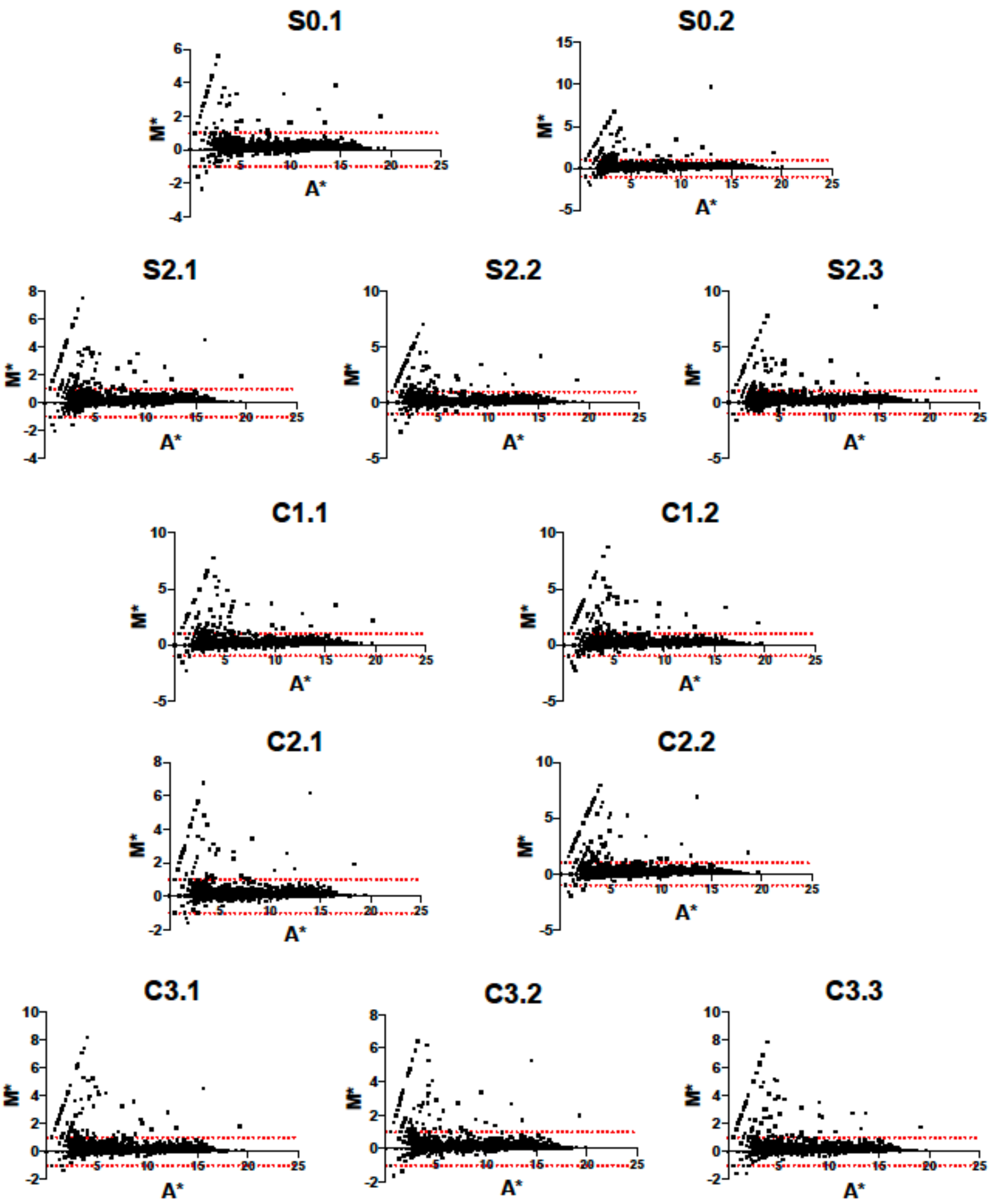


**Figure S3. RNAseq MA* plots of pipeline differences (related to Fig. 2).** Modified MA plots of gene expression results for each sample are shown. M* = log2(1 + Y ) - log2(1 + X) with a red line indicating the threshold for flagging a gene (±1). A* = ½(log2(1 + Y ) + log2(1 + X)).

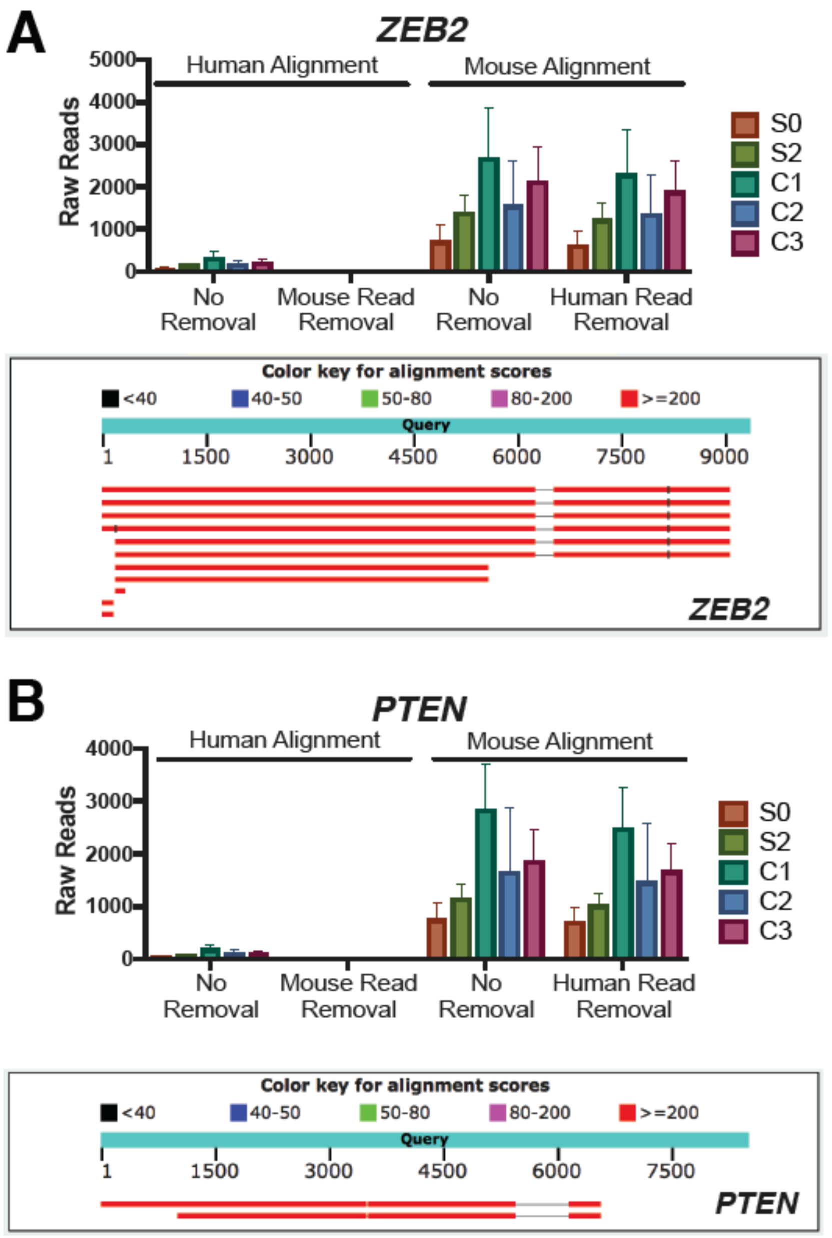


**Figure S4 (related to Fig. 2). *ZEB2* and *PTEN* mRNA homology between mouse and human.** (**A**) (Top) Raw reads mapped to human *ZEB2* before and after mouse alignment read removal versus raw reads mapped to mouse *Zeb2* before and after human alignment read removal. (Bottom) Screenshot of BLAST results (19) for alignment of the human reference *ZEB2* mRNA sequence (NM_014795.4) downloaded from the NCBI database (21) against the mouse RefSeq RNA database. Alignment scores are shown in Table S4. (**B**) Same as above for *PTEN* (NM_000314.7). Error bars represent standard deviation.

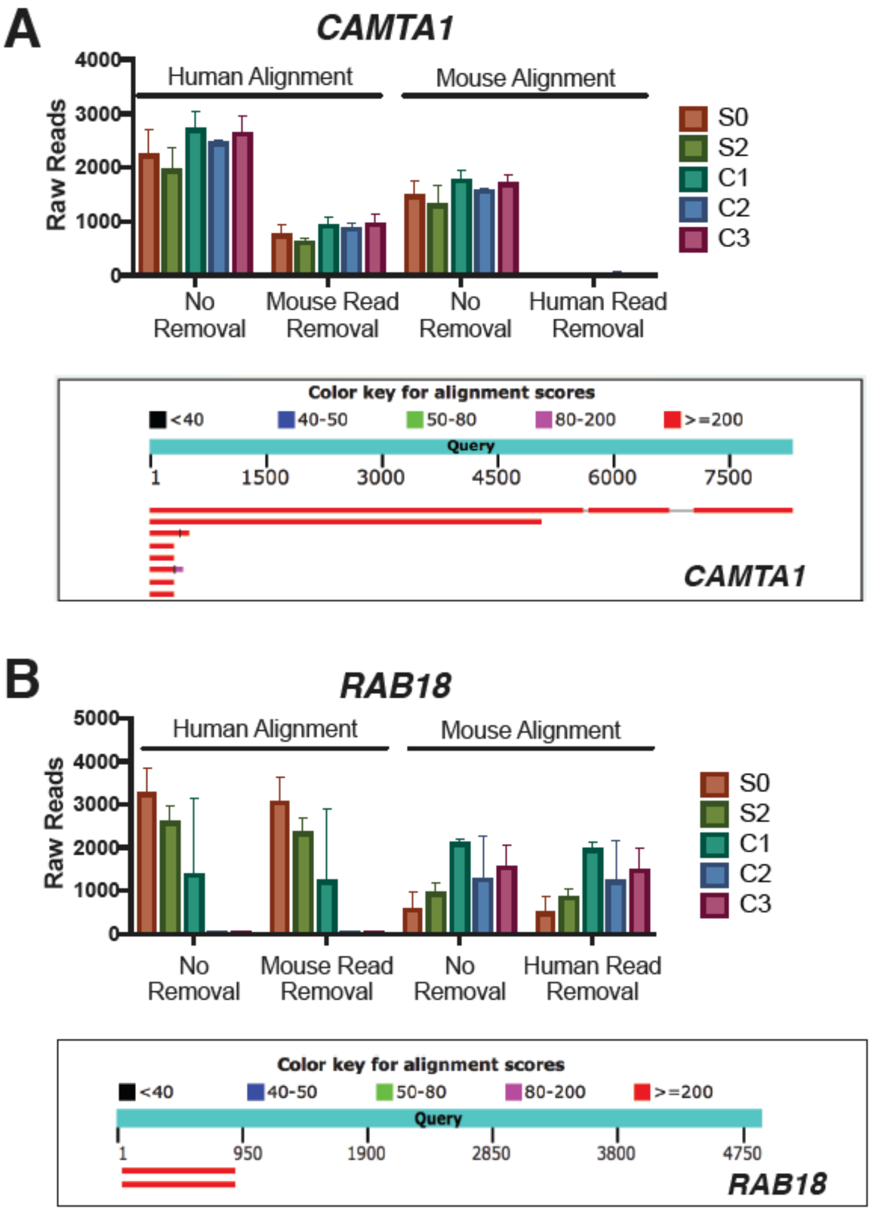


**Figure S5 (related to Fig. 2). *CAMTA1* and *RAB18* mRNA homology between mouse and human.** (**A**) (Top) Raw reads mapped to human *CAMTA1* before and after mouse alignment read removal versus raw reads mapped to mouse *Camta1* before and after human alignment read removal. (Bottom) Screenshot of BLAST results (19) for alignment of the human reference *CAMTA1* mRNA sequence (NM_015215.4) downloaded from the NCBI database (21) against the mouse RefSeq RNA database. Alignment scores are shown in Table S4. (**B**) Same as above for *RAB18* (NM_021252.5). Note that the mouse read removal does not impact the interpretation of *RAB18* as a significantly changed gene in PH039 tumor evolution. Error bars represent standard deviation.

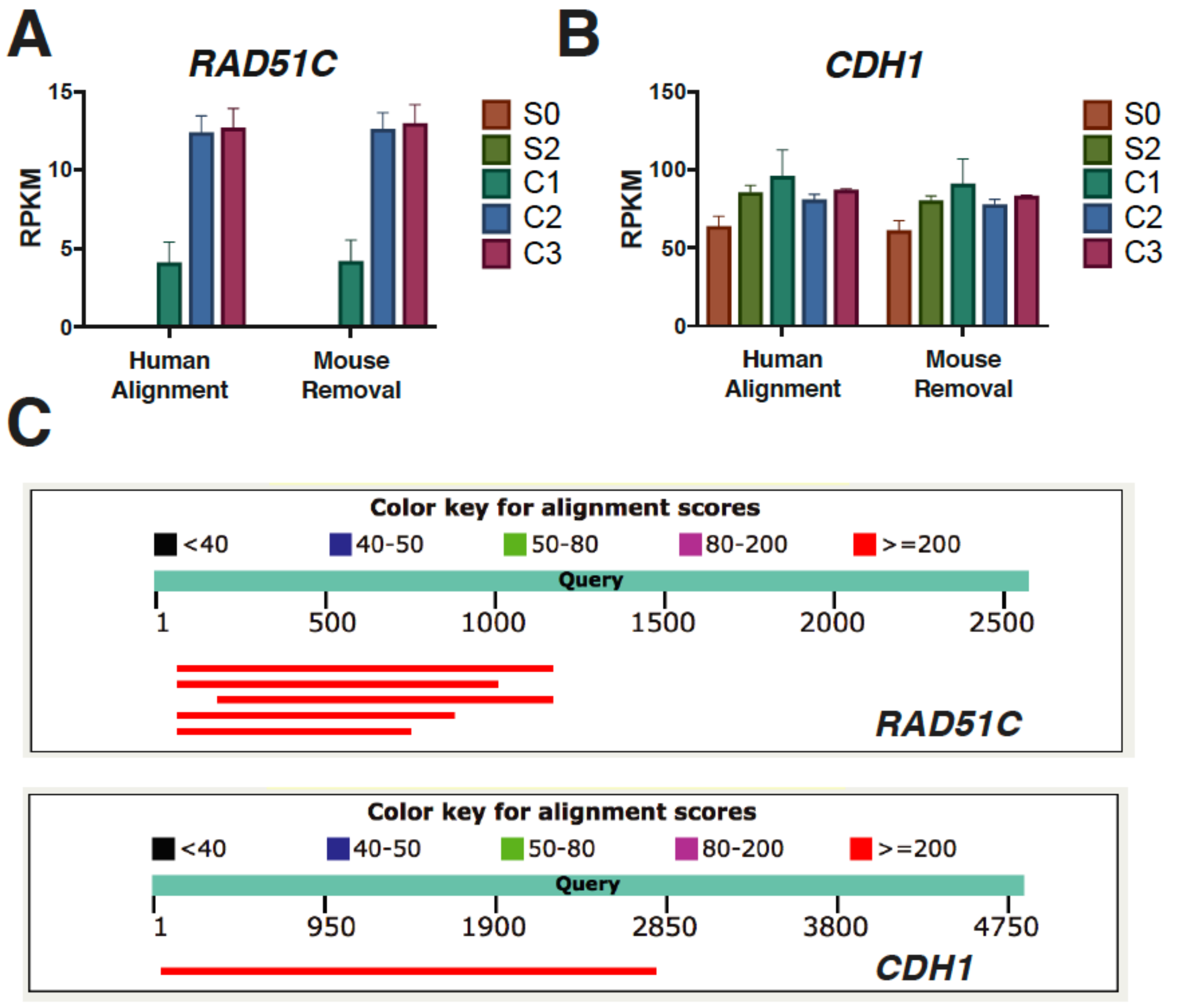


**Figure S6 (related to Fig. 2). *RAD51C* and *CDH1* mRNA homology between mouse and human.** (**A**) (Top) Raw reads mapped to human *RAD51C* before and after mouse alignment read removal versus raw reads mapped to mouse *Rad51c* before and after human alignment read removal. (Bottom) Screenshot of BLAST results (19) for alignment of the human reference *RAD51C* mRNA sequence (NM_058216.3) downloaded from the NCBI database (21) against the mouse RefSeq RNA database. Alignment scores are shown in Table S4. (**B**) Same as above for CDH1 (NM_004360.5). Error bars represent standard deviation.

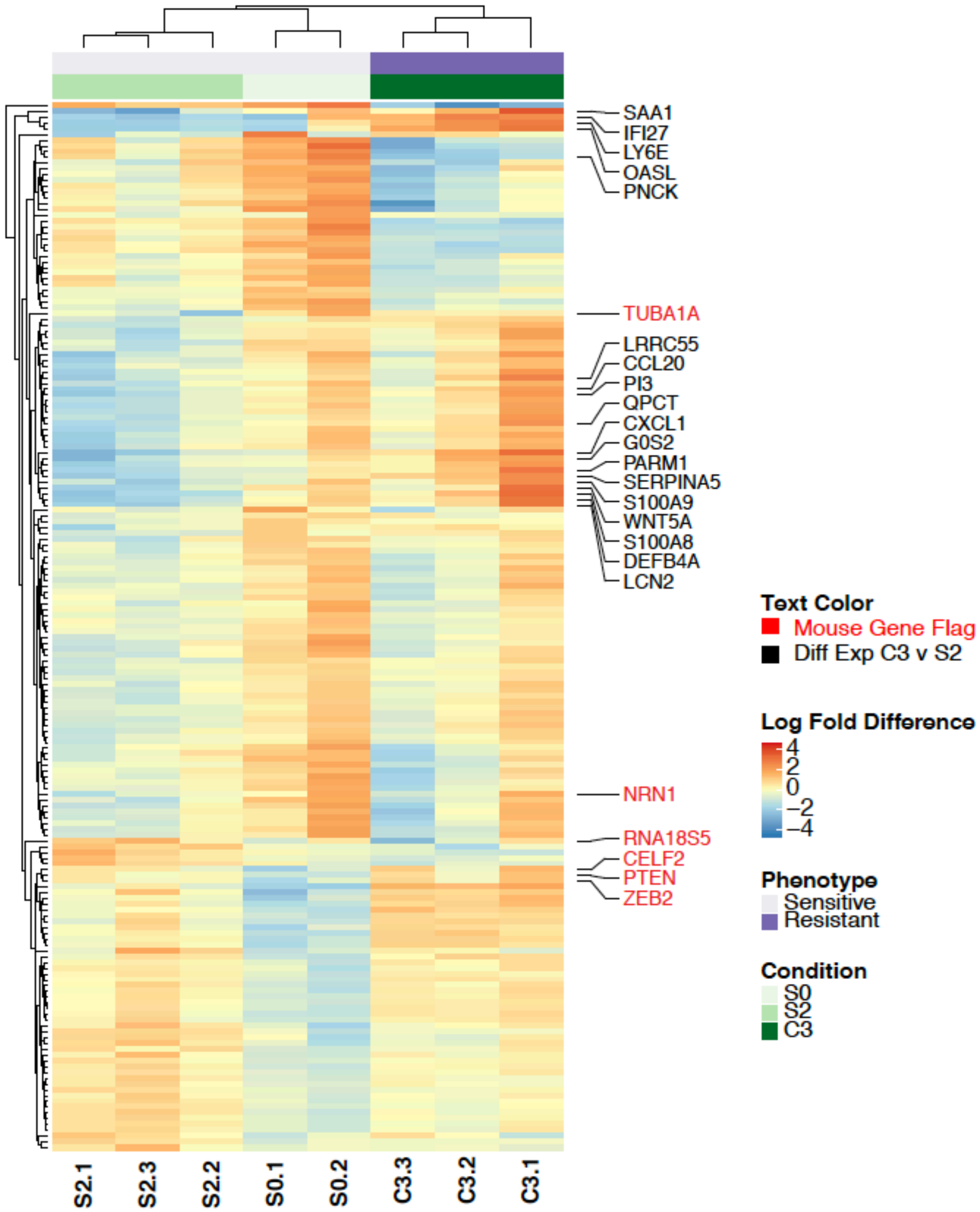


**Figure S7 (related to Fig. 2). Evolution of PH039 signaling pathways through selective passaging.** Pairwise differential expression comparison of S0 and S2 yielded 181 genes with differential expression between S0 and S2 (absolute value of log2 fold change >1 and p-value < 0.01). Raw counts per sample are normalized to counts per million and rows are mean normalized. Labeled genes (black) indicate genes for which the pairwise comparison of S2 and C3 were highly changed (absolute value of log2 fold change >2 and p-value < 0.01). Labeled genes (red) indicate genes which were flagged as differentially expressed between mouse and human based on the criteria outlined in the Supplemental Results section entitled "Considerations of multi-mapping in the analysis of the PDX RNAseq data." Clustering (rows and columns) is unsupervised UPGMA clustering on the displayed genes.

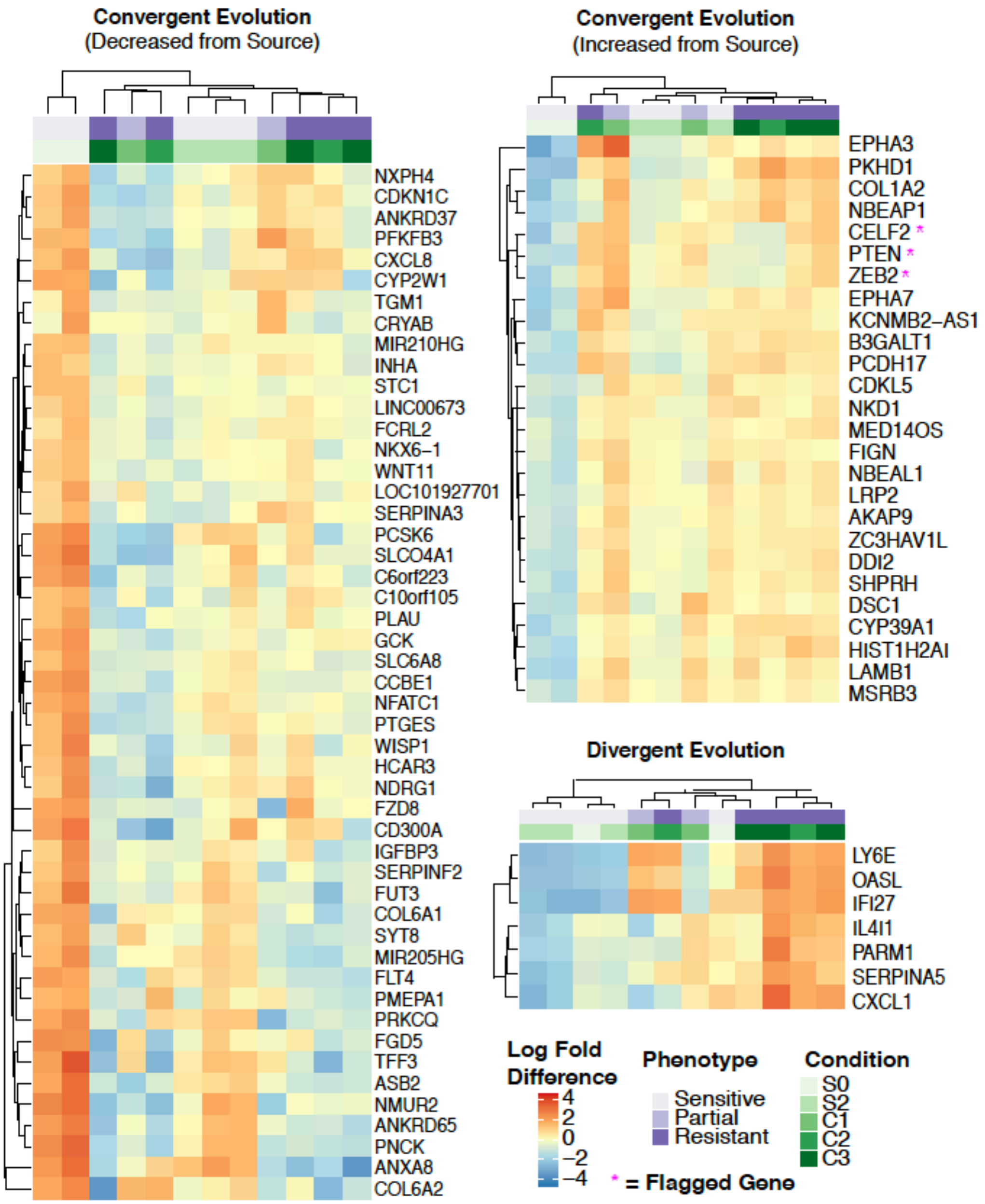


**Figure S8 (related to Fig. 2). Convergent and divergent evolution of genes through selective passaging and drug treatment**. Source tumor (S0) was compared with S2 and C3 to identify increased and decreased genes (absolute value of log2 fold change >1 and p-value < 0.01). Genes that were both up-regulated or down-regulated from source are shown displaying convergent evolution. Genes that diverged in opposite direction from source tissue are shown in the divergent evolution heatmap. Flagged genes are represented with an asterisk and represent those differentially expressed between mouse and human based on the criteria outlined in the Supplemental Results section entitled “Considerations of multi-mapping in the analysis of the PDX RNAseq data.” Raw counts per sample are normalized to counts per million and rows are mean normalized and clustering (rows and columns) is UPGMA clustering.

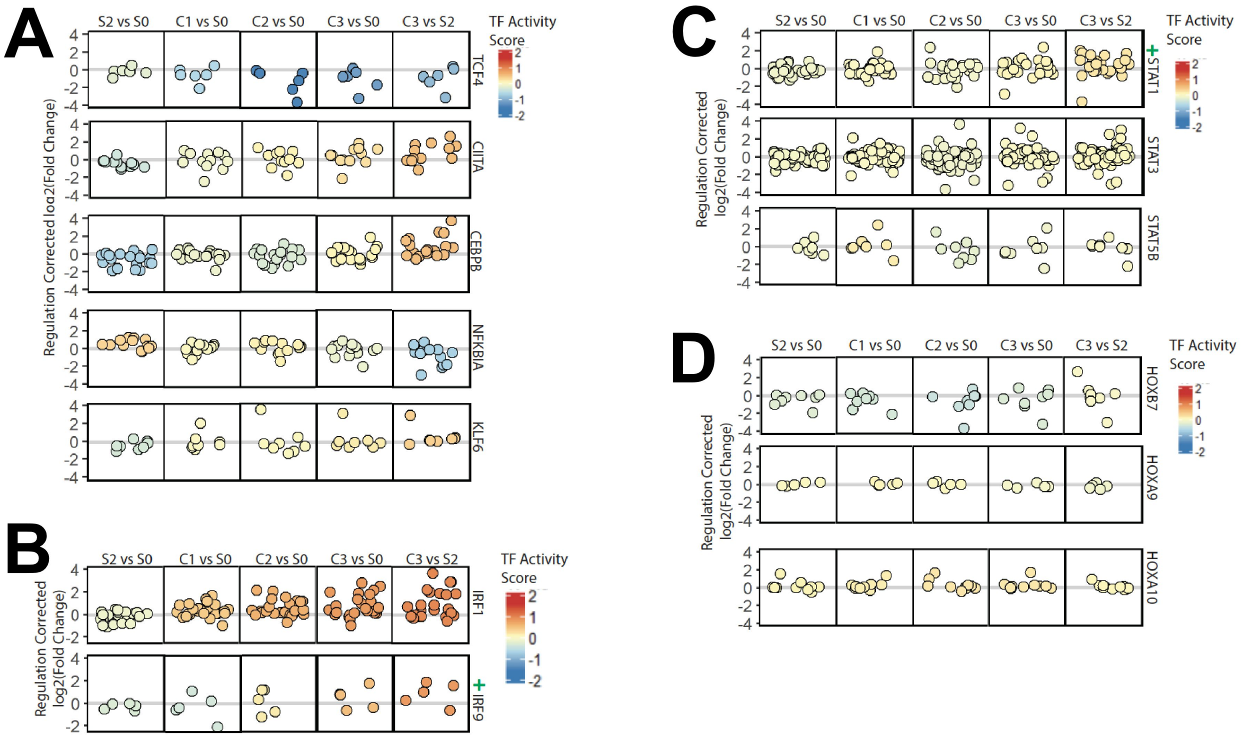


**Figure S9 (related to Figure 2). Additional changes in transcription factor activity score. A,** transcription factors ranked 6-10 for absolute values of changes in activity scores. **B,** changes in activity scores of IRF1 (from Fig. 2B for comparison) and IRF9. **C, D,** changes in STAT and HOX family member activities scores, respectively. Transcription factors with greater than one log2 fold change themselves are indicated by a green plus sign (+) by their row label.

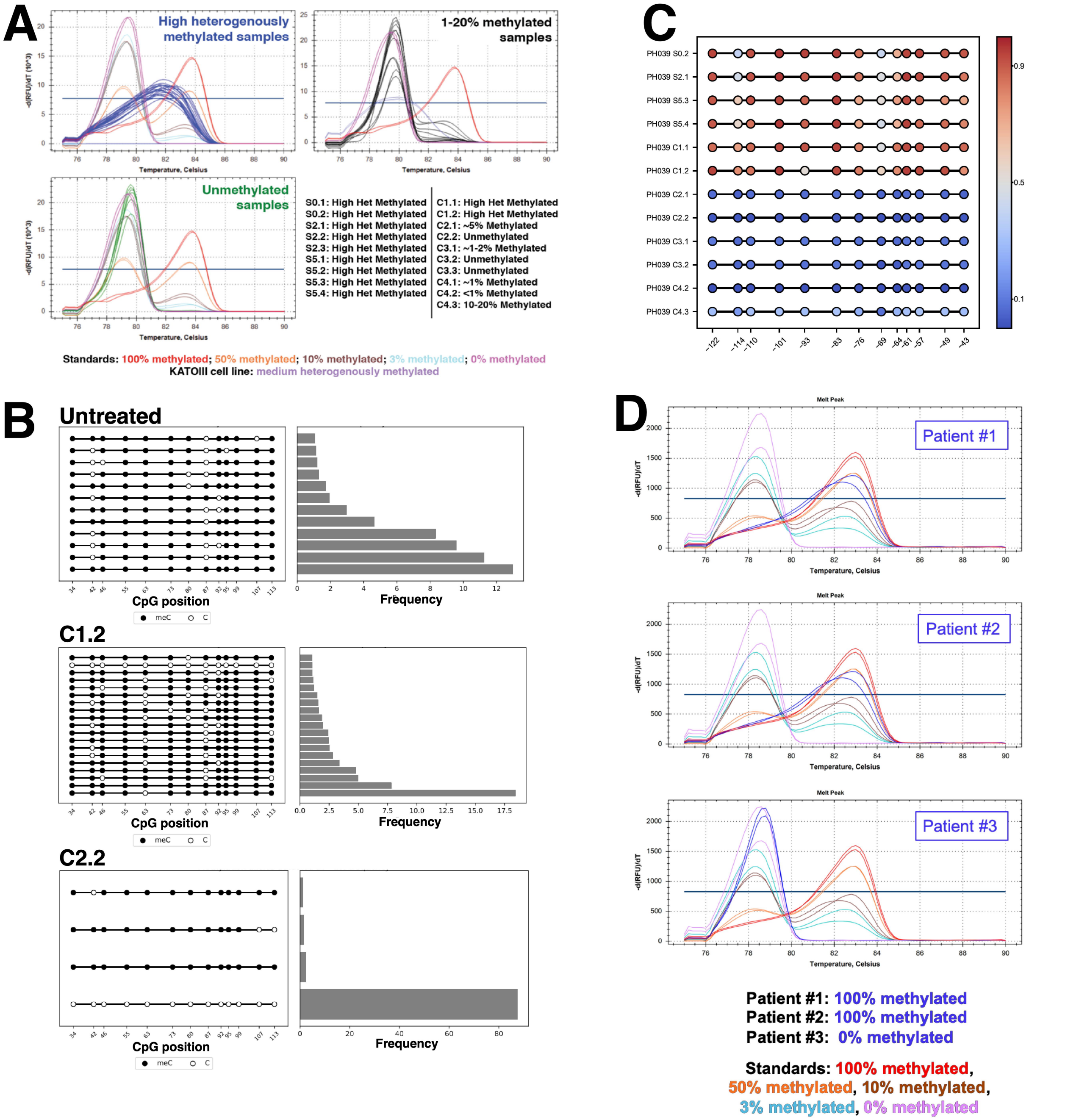


**Figure S10. Further analysis of *RAD51C* methylation loss in PH039 and clinical samples (related to Figure 3). A,** MS-HRM analysis showed samples with heterogeneous methylation, 1-20% methylation, and lack of methylation of the *RAD51C* promoter. Source tissue remained heterogeneously methylated throughout multiple passages. Additionally, the methylation was maintained through one round of niraparib treatment. Following two rounds of treatment, at which time the tissue was resistant (C2), methylation was <20%. Samples were analyzed in one run, with samples split by methylation for visual purposes. All results are summarized by sample ID. Samples and standards are denoted in the color that they appear on the graph. **B,** allele-specific RRBS results for representative samples from four untreated tumors (S0-S5), two tumors analyzed at C1 and two tumors analyzed at C2. Shown for each sample are allele-specific methylation patterns that occurred with >1% frequency in the sample (left) and the frequency of reads with that pattern (right). **C,** frequency of methylation of specific CpGs determined by allele-specific RRBS on DNA from each of the indicated samples. Numbers below panel indicate location relative to the transcriptional start site. **D,** MS-HRM analysis on the samples depicted in Fig. 3H**.**

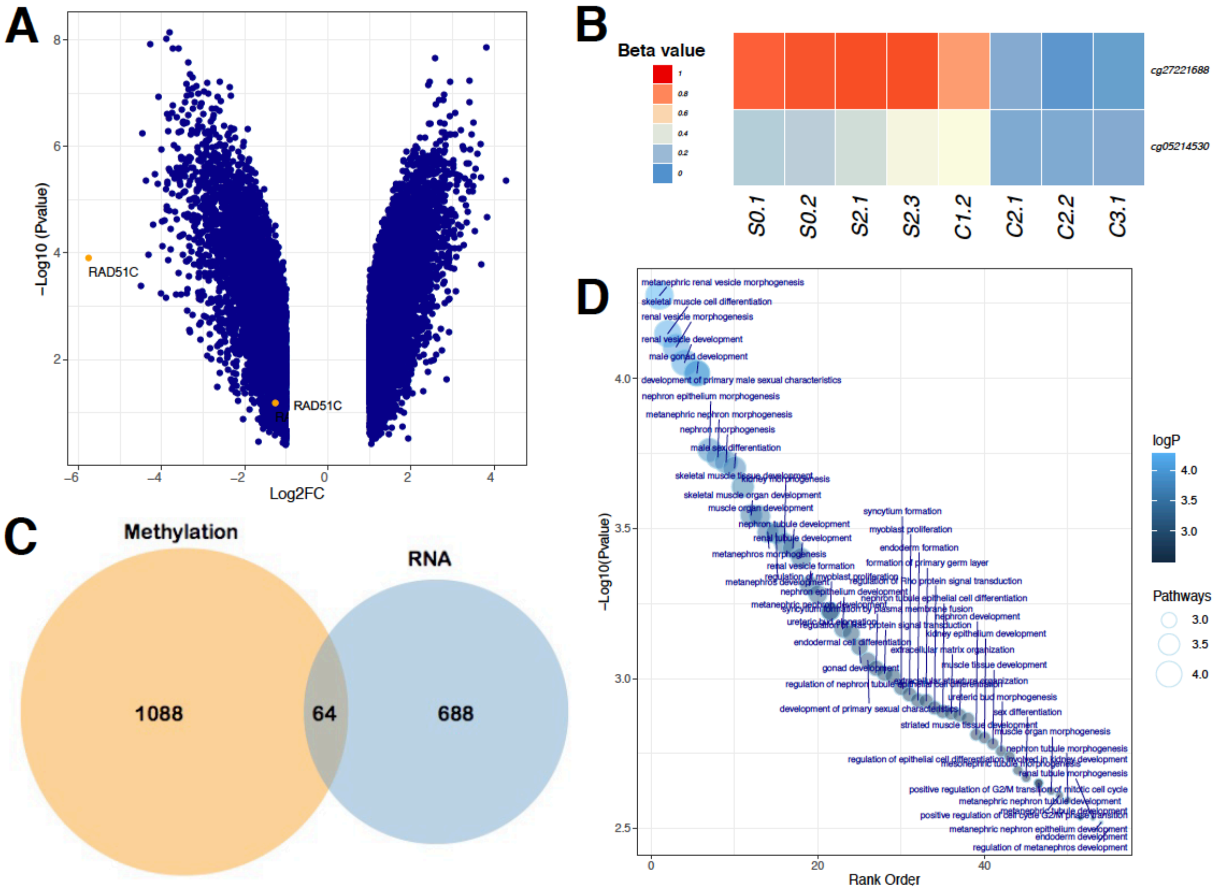


**Figure S11. Changes in DNA methylation assessed by EPIC methylation array (related to Figure 3).** **A,** volcano plot showing EPIC array probes with $log_2FC > 1$. **B,** beta values of the two probes for the *RAD51C* promoter across the specimens harvested at timepoints depicted in Figure 1A. **C,** Venn diagrams showing overlap of changes in methylation (absolute $log_2FC > 2$) and RNA (absolute $log_2FC > 1$, $p < 0.05$). **D,** pathways involving genes that have probes with $log_2FC > 2$.

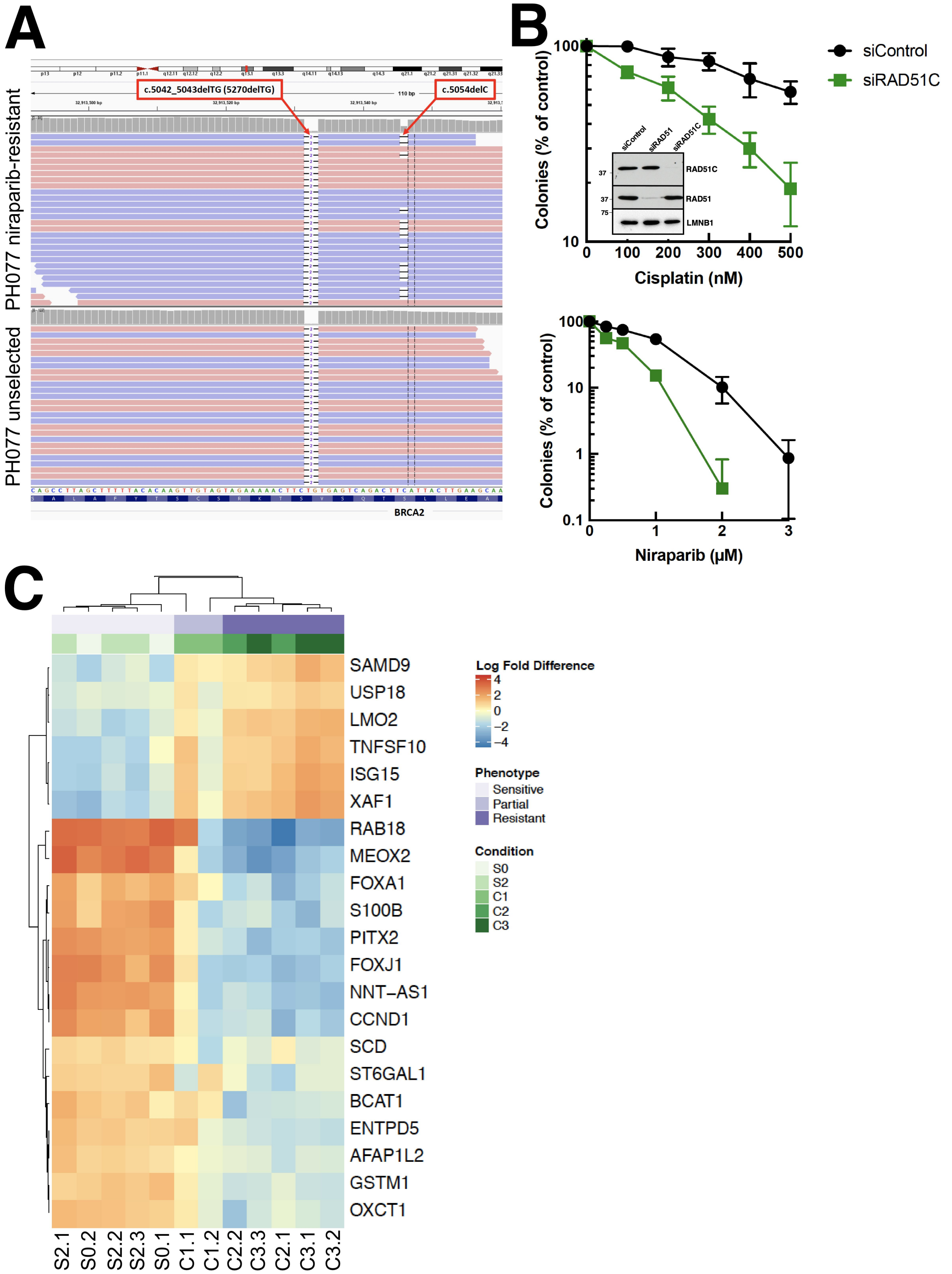


**Figure S12 (related to Fig. 4). A,** IGV screenshot showing original BRCA2 frameshift mutation, c.5042_5043delTG (5270delTG) at ~100% variant allele frequency and secondary reversion mutation c.5054delC in niraparib-selected PDX. **B,** effect of pooled RAD51C siRNA on sensitivity to cisplatin and niraparib. **Inset in B**, RAD51C siRNA knocks down RAD51C but not RAD51. **C,** heatmap showing 21 transcripts that have been implicated in platinum sensitivity. qRT-PCR to confirm a subset of these changes is shown in Fig. 4D.
